# Associations between *in vitro*, *in vivo* and *in silico* cell classes in mouse primary visual cortex

**DOI:** 10.1101/2023.04.17.532851

**Authors:** Yina Wei, Anirban Nandi, Xiaoxuan Jia, Joshua H. Siegle, Daniel Denman, Soo Yeun Lee, Anatoly Buchin, Werner Van Geit, Clayton P. Mosher, Shawn Olsen, Costas A. Anastassiou

## Abstract

The brain consists of many cell classes yet *in vivo* electrophysiology recordings are typically unable to identify and monitor their activity in the behaving animal. Here, we employed a systematic approach to link cellular, multi-modal *in vitro* properties from experiments with *in vivo* recorded units via computational modeling and optotagging experiments. We found two one-channel and six multi-channel clusters in mouse visual cortex with distinct *in vivo* properties in terms of activity, cortical depth, and behavior. We used biophysical models to map the two one- and the six multi-channel clusters to specific *in vitro* classes with unique morphology, excitability and conductance properties that explain their distinct extracellular signatures and functional characteristics. These concepts were tested in ground-truth optotagging experiments with two inhibitory classes unveiling distinct *in vivo* properties. This multi-modal approach presents a powerful way to separate *in vivo* clusters and infer their cellular properties from first principles.

## Introduction

The cellular composition of the brain is diverse with recent studies in rodent neocortex identifying tens of cell types^1–4^. The expectation is that these types serve distinct roles in behavior. However, disentangling their function is challenging. The difficulty is twofold. First, extensive single-cell characterization of neurons, mainly propelled by advances in sequencing technology, allow sampling from large populations at the cellular level, revealing a multitude of cell types. These types exist within detailed, molecular-based taxonomies of neocortex, hippocampus and other brain circuits^3, 5, 6^. *In vitro* cellular electrophysiology and morphology reconstructions, in turn, offer a phenomenology-based approach in defining taxonomies that is easier translated to *in vivo* dynamics, e.g. via spike response properties^7, 8^. Taxonomies accounting for the three main data modalities simultaneously are scarce, with a few noteworthy exceptions^2, 9, 10^.

The second challenge lies in monitoring cell classes identified via their *in vitro* molecular, electrophysiology and morphology properties *in vivo*. *In vivo* imaging of virally or genetically targeted populations offer remarkable insights in how these populations organize during behavior but are unable to resolve single action potentials due to their low sampling rate and the highly nonlinear relationship between spikes and calcium indicator fluorescence^11–13^. Single-wire or high-density extracellular electrophysiology recordings, on the other hand, offer much improved temporal resolution to monitor spiking and spike-related activity *in vivo* even if their ability to resolve cell types is limited. Typically, a handful of spike features can separate between major classes, e.g., the extracellular action potential (EAP) width separates fast-spiking (FS) from other so-called regular-spiking (RS) units^14–17^. Early slice experiments indicated that RS and FS cells probably correspond to pyramidal cells and interneurons, respectively^18^, while other studies found a more intricate correspondence^17, 19, 20^. With recent advancements drastically increasing the electrode density of silicon probes^21^, spatiotemporal information on EAP waveforms increased significantly allowing for more refined clustering of *in vivo* EAPs^22, 23^. Even so, linking cellular taxonomies to *in vivo* signatures, i.e., classification, in a systematic manner for *in vivo* recordings has been difficult.

Single-cell computational models make it possible to link various types of data by incorporating constraints and generating predictions across data modalities, e.g., predicting a particular ion conductance based on properties of the electrophysiological response such as spike shape or frequency. In a recent study, a large-scale model generation and evaluation effort developed bio-realistic, single-cell models for mouse primary visual cortex (V1) accounting for ion conductances along the entire neural morphology^24^. Importantly, these models closely capture distinguishing properties of major excitatory and inhibitory classes integrating electrophysiology, morphology and transcriptomics data. A key aspect of conductance-based models is their ability to emulate extracellular electrophysiology signatures such as the EAP-waveform^25–27^. Thus, these models integrate a variety of data modalities they were trained on (electrophysiology and morphology) or validated against (transcriptomics) and predict a fourth data modality, i.e., the EAP waveform and its associated features.

Here, we show that unsupervised clustering of mouse V1 units recorded via high-density Neuropixels probes ^21^ results in two one-channel and six multi-channel clusters with distinct EAP and EAP-propagation profiles, respectively. Importantly, these clusters exhibit functional differences and distinct coupling to endogenous oscillations, i.e. the main criterion for being considered truly distinct populations in the microcircuit. To determine the differences between the individual clusters we use biophysical models that capture single-cell data from cortical transgenic mouse lines to define EAP templates. Using a supervised classifier, we show that morphological spiny vs. aspiny neurons closely map to RS and FS units, respectively, recorded *in vivo*. Next, we map the six multi-channel clusters with their distinct EAP propagation profiles to model populations, compare between model population setups and identify conductances and morphology features that explain the EAP differences between the *in vivo* clusters. Our newfound ability to separate between clusters is exemplified in ground-truth, optotagging experiments where we separate between two major inhibitory classes *in vivo* and show their distinct entrainment profile to ongoing neocortical oscillations.

## Results

### Extracellular action potential recordings from *in vivo* experiments and biophysical models of cell types

Analysis of extracellular action potential (EAP) waveforms of so-called “units” (putative single neurons) typically clusters into two groups, regular-spiking (RS) vs. fast-spiking (FS) (**Fig. 1a**). We sought a more refined classification scheme using data from a recent *in vivo* survey of electrophysiological activity in awake mice^23^. We focused on data from units in primary visual cortex (V1) recorded using Neuropixels probes (**Fig. 1b**). These probes offer a dense arrangement of recording sites (**Fig. 1c**), which allows EAP signals from single units to be detected on multiple recording channels (**Fig. 1c**; example unit #1: a FS unit; example unit #2: a RS unit; bold: channels with largest EAP amplitude). We analyzed units from 25 wild-type mice, 8 mice expressed ChR2 in parvalbumin-positive cells (Pvalb), and 12 in somatostatin-positive cells (Sst) (**Fig. 1d**). We only analyzed units located in V1 with an average of 48 units per wild-type mouse being well-isolated (unit isolation criteria: see Methods; **Fig. 1d**; total number of units = 1204) during spontaneous activity. The depth of layer 4 was determined from where the visual stimulus (flash) evoked a strong response in the current source density (CSD) ^28, 29^ (**Fig. S1**). Unit location along the cortical depth was adjusted relative to layer 4 (depth 0 indicates the center of layer 4). The estimated soma location of well-isolated units (based on EAP properties) in our study spanned from layers 2/3 through 6 with the majority located in layers 4 and 5 (**Fig. 1d**).

**Figure 1.**
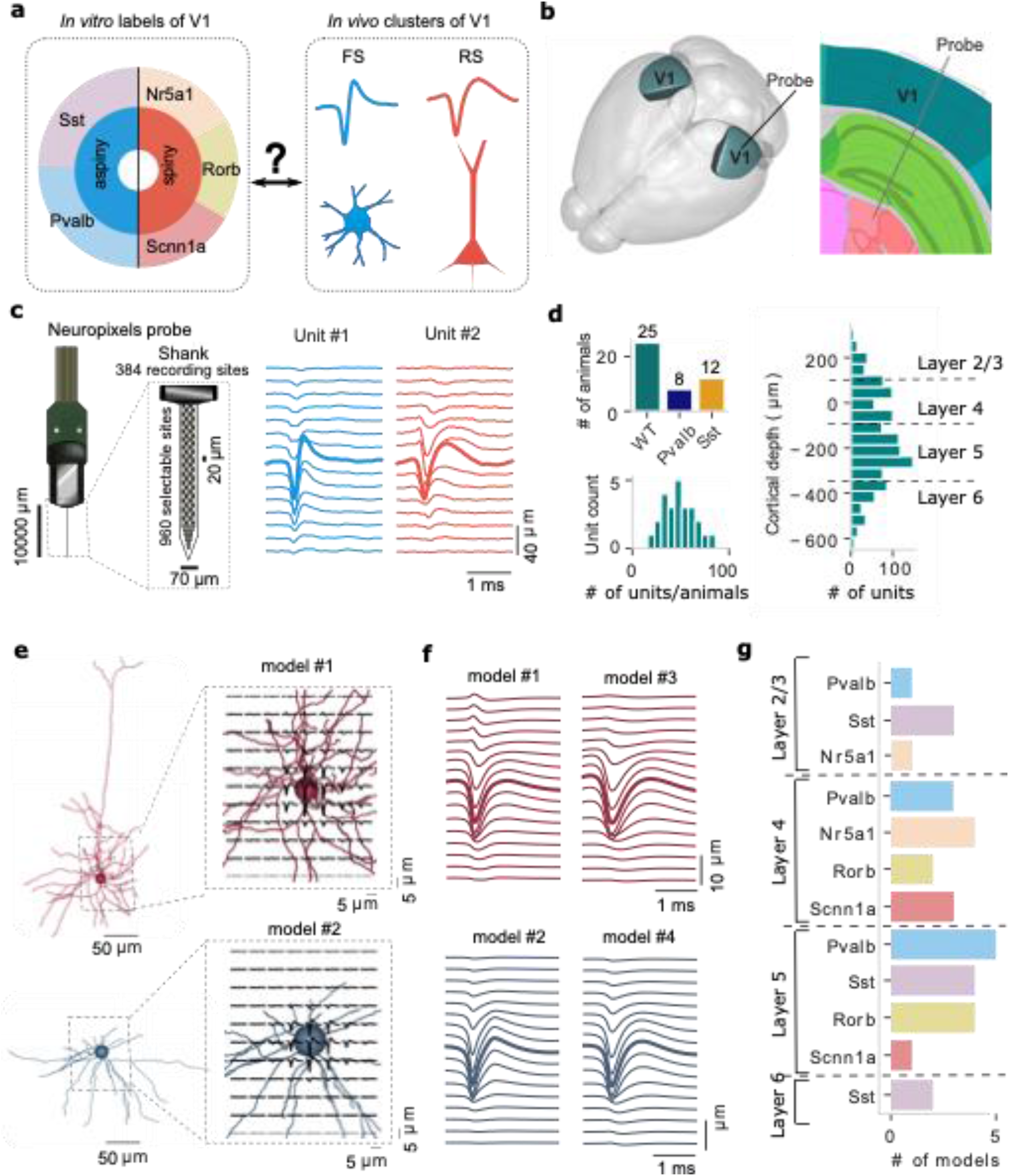
Extracellular action potential (EAP) recordings from *in vivo* experiments (a-d) and single-cell modeling (e-g). **a**) Left, labels for Cre-line and morphology (spiny vs. aspiny) groups of single neurons used in this study characterized *in vitro* via intracellular electrophysiology and morphology reconstructions. Right, *in vivo* EAP waveform analysis typically results in two clusters, fast-spiking (FS) vs. regular-spiking (RS) units. **b**) Primary visual cortex (V1) in the mouse brain (left) and typical cortical depth placement of a Neuropixels probe along V1. **c**) The 384 electrode sites of the Neuropixels probe are densely arranged along the linear shank probe (left; 20 μm vertical spacing, 2 sites per row; black squares: location of recording sites). EAP waveforms from two example units (unit #1: FS; unit #2: RS) including the channel with the largest amplitude (closest to the soma, bolded lines) and channels above and below the soma. **d**) Top: the number of Neuropixels-implanted mice for wild-type (n=24), parvalbumin-expressing (Pvalb, n=8) and somatostatin-expressing (Sst, n=12); bottom: the distribution of units per wild-type mouse recorded in V1 during drifting gratings (total number of units = 1204). Distribution of units along the V1 depth axis with 0 indicating the center of layer 4. **e**) Bio-realistic, single-cell models of V1 (“all-active”) are generated from *in vitro* experiments and activated via synaptic background to elicit intracellular activity and associated EAP signals in the vicinity of the cellular morphology. The cellular morphology is represented with a spherical soma and full dendritic reconstruction (axon not shown). Example simulations of EAP signals are shown for a spiny (top: red, cell ID: 395830185) and an aspiny (bottom: blue, cell ID: 469610831) single-cell model. **f**) Four examples of the multi-channel EAP including the channel with the largest amplitude (bolded lines, closest to the soma) and channels above and below the soma (top: 2 spiny models; bottom: 2 aspiny models). **g**) In total, 33 single-cell models (15 spiny and 18 aspiny) were generated using a computational optimization framework and included in the study covering a range of major reporter lines and cortical depths. Source data are provided as a Source Data file.

To map the recorded EAP waveforms to specific cell classes we used biophysically detailed models of single neurons. These biophysical models are developed in an unsupervised manner using a multi-objective optimization platform that relies on standardized electrophysiology features and the reconstructed cellular morphology to distribute a set of ionic conductances relevant for cortical neurons^24^. We developed single-cell models that represent a diverse set of transgenic mouse lines to ensure broad coverage across cortical layers and classes^1^. For our study we accounted for 15 spiny (SP) and 18 aspiny (AP) single-cell, so-called “all-active”, biophysically realistic models from V1 optimized based on *in vitro* single-cell electrophysiology and morphology (**Fig. S2**). Notably, the SP vs. AP designation in our study is morphology-based and does not reflect any electrophysiology features such as action potential waveform or spike pattern. The experimental data to produce the single-cell models were part of a systematic characterization of mouse visual cortex where a uniform experimental protocol was used to establish a taxonomy based on cellular electrophysiology and morphology^1, 24^.

Beyond reflecting key properties of various cell types in terms of electrophysiology, morphology and transcriptomics^24, 30^, these biophysical single-cell models reproduce EAP signals in the vicinity of the cellular morphology (**Fig. 1e**, top: spiny cell, cell ID: 395830185; bottom: aspiny cell, cell ID: 469610831). Our computational approach simulated the recording sites of a Neuropixels probe (see Methods; ^27^) resulting in signals emulating *in vivo* unit recordings (**Fig. 1f**). In total, 15 spiny (Cre-reporter lines: 5 Nr5a1, 4 Scnn1a, 6 Rorb) and 18 aspiny (Cre-reporter lines: 9 Pvalb, 9 Sst) single-cell models were developed and included in the study covering a range of major reporter lines and cortical depths (**Fig. 1g**) and especially layers 4 and 5 in accordance with the *in vivo* experiments (**Fig. 1d**).

### The standard waveform features reveal two clusters: RS and FS

Spontaneous and visually evoked activity (flashes) is recorded *in vivo* in head-fixed animals implanted with Neuropixels probes in V1 while running freely on a rotating disc (**Fig. 2a**; *N* = 1204 units from 25 mice during spontaneous activity). For the EAP analysis, we derived the ***one-channel EAP*** from the channel with the maximum EAP-amplitude (**Fig. 2b**, middle: red bolded trace), while the ***multi-channel EAP*** includes additional channels above and below the maximum EAP channel (**Fig. 2b**, middle). We define two one-channel EAP features (**Fig. 2b**, left): trough-to-peak width (TPW) and repolarization time (REP). TPW measures the time from the EAP trough until the peak. REP measures the time from EAP peak to the half-peak^17, 27, 31^. TPW and REP are usually sufficient to classify units between narrow and wide waveforms^15, 27^, the result of the bimodal distribution of TPW in cortex (**Fig. S3**). We also found two major clusters in our *in vivo* data, i.e. a narrow TPW cluster with reduced REP (**Fig. 2d**, bottom, blue) and a wide TPW cluster of increased REP (**Fig. 2d**, bottom, red), respectively. Both the elbow method and density method of unsupervised *K*-means clustering^22^ independently confirmed the optimal number of clusters are two. Specifically, the narrow waveform units exhibit lower TPW (**Fig. S3**) and lower REP (**Fig. S3**) than the wide waveform units. Furthermore, narrow waveform units (n=281, 23.3%) exhibit elevated spike frequency vs. wide waveform ones (n=923, 76.7%): narrow waveform units spike at a median firing rate of 4.85 Hz (interquartile range, IQR: 1.93-10.79 Hz) while wide waveform units fire at median of 2.05 Hz (IQR: 0.84-5.00 Hz). Thus, narrow waveform units spike significantly faster than their wide waveform counterparts (Mann-Whitney U test, p=3.5*10^-18^; **Fig. S3**). We conclude that narrow EAP waveforms approximately map to fast-spiking (FS) units while wide waveforms approximately correspond to regular-spiking (RS) units (Supplementary Data 1).

**Figure 2.**
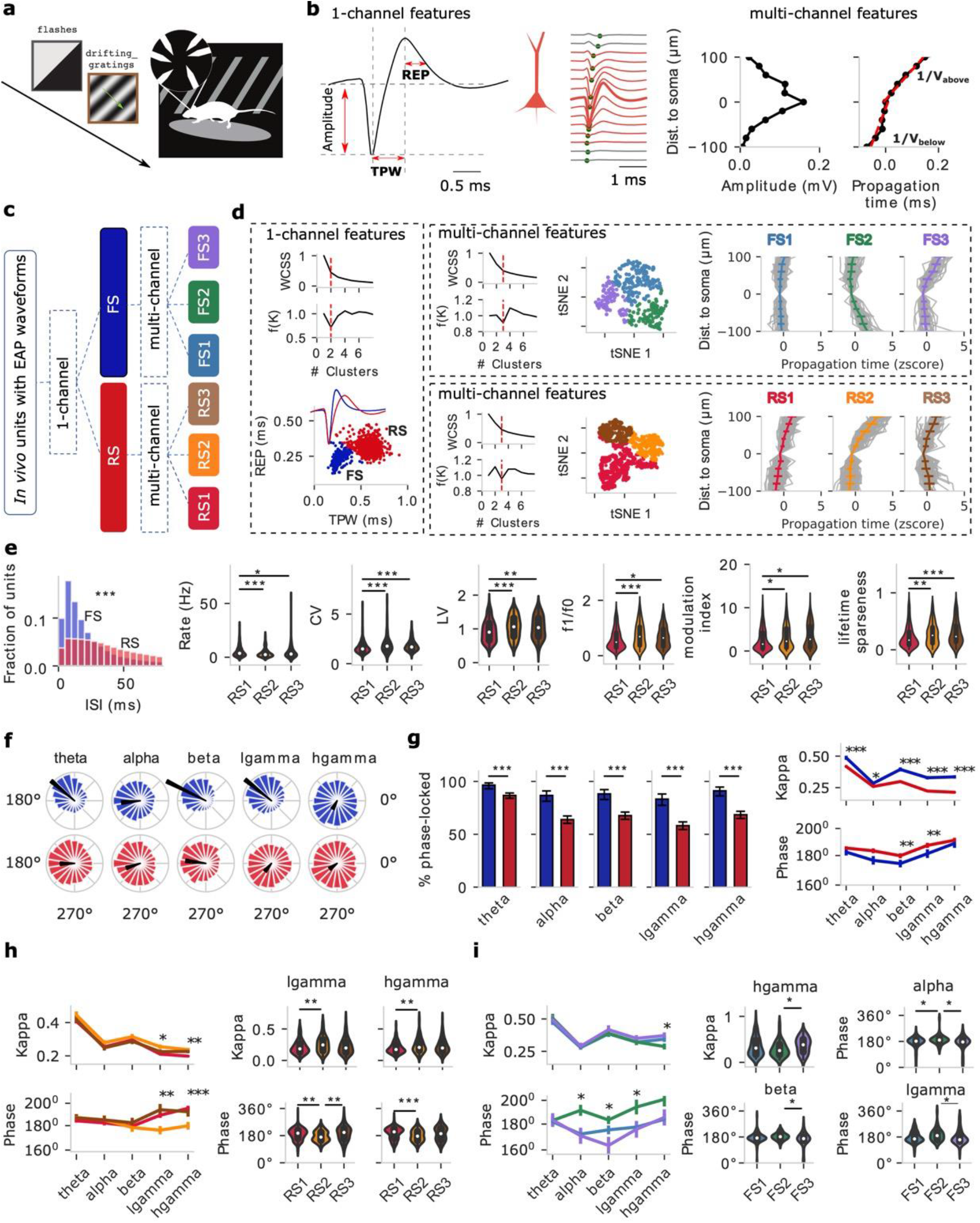
Clustering of *in vivo* V1 units from wild-type mice based on extracellular action potential (EAP) features during drifting gratings results in two one-channel and six multi-channel clusters with distinct EAP properties. **a)** Animals are exposed to visual stimuli (e.g., flashes, drifting gratings) while running on a wheel with Neuropixels probes recording extracellular V1 activity. **b)** One-channel EAP waveform features (left) from the location of the largest EAP amplitude: trough-peak width (TPW), and repolarization time (REP). Multi-channel EAP waveform features: the inverse of propagation velocity below (1/*V_below_*) and above (1/*V_above_*) soma are separately estimated by linear regression (right, red lines). **c)** Unsupervised clustering on one-channel EAP features (TPW and REP) results in two major populations, fast-spiking (FS) and regular-spiking (RS) units. Subsequently, unsupervised clustering of each population using multi-channel EAP features (1/*V_below_* and 1/*V_above_*) results in three clusters, respectively. c) One-vs. multi-channel clusters. **d)** The one-channel clusters (923 RS and 281 FS from 25 wild-type mice, left), multi-channel clusters FS1-3 (right top) and multi-channel clusters RS1-3 (right bottom) are shown including two clustering metrics: within cluster sum of squares (WCSS) and density function. The red dotted line indicates the number of optimal clusters. t-distributed stochastic neighbor embedding (t-SNE) for FS1-3 (right top, n=130 FS1, n=82 FS2, n=69 FS3) and RS1-3 (right bottom, n=479 RS1, n=235 RS2, n=209 RS3) units based on features extracted from multi-channel waveforms. The spatial propagation of EAPs is distinct for the clusters (gray: individual units). Data are presented as mean ± SD (standard deviation). e) One-channel FS and RS clusters show distinct interspike interval (ISI) distributions (Mann-Whitney U test, two-sided, p=0.0, total 2900284 spikes of FS, 4586637 spikes of RS). Response properties of the multi-channel clusters to drifting gratings shows that RS1-3 exhibit distinct properties in the overall excitability (spike rate, coefficient of variation: CV, local variation: LV, n=419 RS1, n=173 RS2, n=153 RS3) and stimulus-dependent response characteristics (f1/f0, modulation index, lifetime sparseness, n=430 RS1, n=182 RS2, n=156 RS3). Kruskal-Wallis H-test; p-values corrected using the Holm-Bonferroni method for multiple tests. *p<0.05, **p<0.01, ***p<0.001. f) Phase distribution of examples of a FS (blue) and RS (red) unit at theta, alpha, beta, low gamma (lgamma), high gamma (hgamma) frequency band (black arrow: preferred phase and kappa). g) Left: The percentage of phase-locked units of one-channel FS (n=203) and RS (n=745) clusters at different LFP frequency bands: theta, alpha, beta, low gamma, high gamma. Two sample z test for proportions with p values corrected by the Holm-Bonferroni method for multiple tests. Right: kappa and preferred phase. Data are presented as mean ± SEM (standard error of mean). Mann-Whitney U test, two-sided, *p<0.05, **p<0.01, ***p<0.001. h-i) Phase-locking analysis of multi-channel RS (d, n=419 RS1, n=173 RS2, n=153 RS3) and FS (e, n=100 FS1, n=54 FS2, n=49 FS3) clusters to ongoing oscillations in different LFP bands. Kruskal-Wallis H-test; p-values corrected using the Holm-Bonferroni method for multiple tests. *p<0.05, **p<0.01, ***p<0.001. Source data are provided as a Source Data file.

### Spatial features reveal six distinct sub-clusters in mouse V1: 3 RS and 3 FS

Multi-channel EAP waveforms introduce an additional dimension, space, into the analysis. We accounted for the EAP amplitude and the EAP propagation with respect to time (**Fig.2b**, right) as a function of recording distance to the largest EAP location, assumed to be closest to the soma/axon initial segment^25^. For the multi-channel analysis (**Fig. 2b**, middle), we calculate two additional spatial EAP features (**Fig. 2b**, right): the inverse of the EAP propagation velocity below (1/*V_below_*) and above (1/*V_above_*) the soma^22^. 1/*V_below_* and 1/*V_above_* are separately estimated via linear regression (**Fig 2b**, right, red lines). We define a propagation symmetry index, the ratio of 1/*V_below_* and 1/*V_above_*, with a larger symmetry index indicating a more asymmetric propagation, for example, due to the presence of apical dendrites in excitatory pyramidal neurons^32^ . Looking at the multi-channel EAP features of FS vs. RS, RS generally exhibits a more asymmetric EAP propagation below vs. above the putative soma location than FS, **Fig. 2c-d**; **Fig. S3c-d**, middle). We conclude that one-channel clusters RS and FS do not only separate via TPW but, in fact, are also distinct in how their spikes propagate along the extracellular space.

We wondered whether multi-channel EAP features can further inform on the composition of FS and RS. To do so, we adopted the one-channel clusters RS and FS and for each of them employed unsupervised clustering using multi-channel features (1/*V_below_* and 1/*V_above_*) to further subdivide into multi-channel clusters. Unsupervised clustering (*K*-means) indicated that the optimal number of multi-channel clusters within FS and RS is three for each (**Fig. 2d**, right top; cluster # independently estimated by the elbow method and density function). The six groups (FS1-3, RS1-3) exhibit distinct multi-channel signatures. For the RS group, RS1 and RS2 show mostly asymmetric propagation with their main difference being the supragranular propagation velocity, i.e. *V_above_*(RS1) > *V_above_*(RS2) (**Fig. S4**). RS1-3 exhibit significant differences in terms of their spatial spread (Kruskal-Wallis H-test; p-values corrected using the Holm-Bonferroni method for multiple tests), with the EAP propagation of RS3 being more spatially confined than RS1-2 while also exhibiting a faster infragranular spike propagation velocity *V_below_* (**Fig. S4;** Supplementary Data 1). FS1-3 also exhibit distinct propagation signatures: while the propagation profile for FS1 is symmetric and fast above and below the spike initiation location, FS2 and FS3 exhibit an asymmetric and slower, direction-dependent profile. Despite their different propagation profiles, FS1-3 exhibit no significant difference in spatial spread (**Fig. S4**). Looking at the distribution of the cortical depth, the six clusters are distributed differently across V1 layers (**Fig. S5**). We conclude that expanding the set from one-to multi-channel EAP features results in further separation within the RS and FS groups into six finer but distinct groupings, three FS (FS1, FS2, FS3) and three RS (RS1, RS2, RS3) clusters, that spread along the V1 depth axis.

### Distinct functional properties of the *in vivo* clusters

To what extent do the *in vivo* clusters separated by their EAP properties also constitute functionally distinct cell populations? We looked into the *in vivo* dynamics during behavior and whether the six clusters show distinct firing properties during a visual stimulation task (drifting gratings). Inter-spike interval (ISI) analysis shows that multi-channel RS clusters exhibit significantly different firing properties: the ISI median of FS is 19.63 ms with 95% confidence interval (CI) at [19.60,19.67] ms, while the ISI median of RS is 53.37 ms with 95% CI at [53.30,53.43] ms (**Fig. 2e**, Mann-Whitney U test, p=0.0). To assess the temporal structure of spiking during the task we also calculated the coefficient of variation (CV) that measures the variance of ISIs and the local variation (LV) measuring variation in adjacent ISIs. We found that the pattern of RS1 spiking is significantly different compared to RS2 and RS3. Specifically, RS1 units exhibit faster, more stereotyped and less variable spiking than RS2-3 units. RS2-3 units, in turn, exhibit relatively slower and more variable spiking dynamics (**Fig. 2e**). Notably, multi-channel RS clusters also exhibit differences in their response to visual presentation. Several measures that assess visual response properties were calculated (see Methods) and we highlight three relevant for drifting gratings: f1/f0, the modulation index, and lifetime sparseness (**Fig. 2e**). Statistically significant differences emerge between RS1 vs. RS2-3 in terms of the response metrics with RS2-3 exhibiting higher response sensitivity and selectivity over RS1, in agreement with the higher CV and LV seen for RS2-3. No significant difference in terms of visual responses was observed between FS1-3. In summary, we found that RS is composed of functionally distinct clusters that beyond their distinct multi-channel properties also exhibit differences in their *in vivo* activity also during visual behavior.

Another measure to identify functionally distinct populations looks at distinct spike phase-locking to ongoing local field potential (LFP) oscillations^27, 33, 34^. We used the Hilbert transform of the bandpass-filtered LFP to assign each spike an instantaneous phase (**Fig. S6a**) in several frequency bands (theta: 3-8Hz, alpha: 8-12.5Hz, beta: 12.5-30Hz, low gamma: 30-50Hz, high gamma: 50-90Hz; Supplementary Data 2). Starting with one-channel clusters, we found that units exhibit a diverse level of entrainment to the LFP bands (per the Rayleigh test for non-uniformity, see Methods, **Fig. S6b,** **Fig. 2f**) with FS containing a significantly higher percentage of phase-locked units than RS across frequency bands (**Fig. 2f, left**). Notably, FS and RS coupling to *in vivo* oscillations is input- and behavior-dependent, with a much lower percentage of phase-locked neurons detected during spontaneous activity (**Fig. S7**) than during drifting gratings (**Fig. 2f, left**) across frequency bands, an observation in line with other studies (e.g.^13^) In general, the percentage of significantly entrained FS units was high and remained broadly unaffected by the specific LFP bands. In contrast, RS couple preferentially to slow LFP oscillations (theta) with the percentage decreasing for higher frequencies (beta, gamma and high gamma). Pairwise comparison revealed that FS have stronger phase-locking across frequency bands and spike earlier in the cycle than RS for beta and low gamma (**Fig.2f-g,** p-values corrected for multiple tests by Holm-Bonferroni method) in line with neocortical patterns seen in monkey and human^35^, but in contrast with hippocampal oscillations where putative excitatory neurons typically fire earlier than putative inhibitory ones^36^. We conclude that one-channel RS and FS show distinct coupling properties to neocortical oscillation with FS coupling being stronger across bands and FS units firing earlier than RS.

Next, we looked at multi-channel clusters and their dynamics during oscillations. We found significant differences in LFP coupling for RS1-3 in the low and high gamma bands, with RS2 exhibiting stronger phase locking to low and high gamma than RS1 (**Fig. 2h**). The preferred phase of RS1-3 remains similar at 180^0-2000^ (RS2 just below 180^0^ vs. RS1 and RS3 just above 180^0^) (**Fig. 2h**). Cluster-specific entrainment to LFP oscillations is also observed in FS clusters (FS1-3). Specifically, FS3 exhibit stronger phase locking to high gamma than FS2, with distinct preferred phases among the three clusters in alpha, beta and low gamma (**Fig. 2i**). We conclude that in addition to their distinct spiking characteristics, multi-channel clusters exhibit distinct coupling properties to LFP oscillations that depend on the behavior.

We also looked at how spike dynamics and coupling to oscillations changes with cortical depth. Based on distance from pia we defined three regions: supragranular (broadly cortical layers 2-3), granular (cortical layer 4) and infragranular (broadly cortical layers 5-6). Looking at one-channel clusters, FS show consistently stronger phase-coupling than RS across the cortical depth for all LFP bands (**Fig. S6c)**. Interestingly, both FS and RS show strong coupling in theta and beta but a strong reduction in coupling in the intermediate alpha band. This pattern is particularly pronounced in the supragranular and granular regions while in the infragranular region there is reduced coupling, especially for FS, compared to the rest of the cortical depth regions (**Fig. S6c)**. We also note the strong coupling of FS units to high frequency oscillations (e.g. high gamma) especially in the supragranular and granular region, a characteristic of electrotonically compact neurons able to follow very fast synaptic drive. In terms of spike phase, RS and FS spike broadly around the same phase with the exception of the granular region where significant differences emerged between FS and RS for beta and gamma bands. Looking at the multi-channel clusters across cortical depth, we found the most significant differences in the coupling strength of RS1-3 in supragranular beta and low gamma with kappa almost doubling between supragranular RS3 and RS1 in beta (**Fig. S6d**). Such diversity in coupling strength among clusters is not observed in granular and infragranular regions though we do find differences in the preferred spike phase of RS1-3 in infragranular layers (**Fig. S6d**). It follows that these multi-channel clusters, except for their distinct multi-channel signatures, also have distinct patterns and role in how they support ongoing cortical oscillations. We conclude that, one-channel RS and FS clusters as well as RS1-3 show distinct coupling patterns along the cortical axis, especially supragranular RS1-3 in the beta bands and infragranular RS1-3 in the gamma bands.

### Multimodal mapping between electrophysiology-, morphology- and Cre-reporter-based classes

What is the cellular identity of the clusters exhibiting such distinct EAP-waveform and *in vivo* properties? To bridge between the *in vivo* clusters and *in vitro* cell classes, we use biophysically realistic single-neuron models of 18 morphologically aspiny (AP) and 15 spiny (SP) mouse neurons (**Table S1**) that capture within cell type variability. These models were generated from two data modalities: the reconstructed morphology and the somatic electrophysiology response resulting from *in vitro* whole-cell patch-clamp experiments^24^. We use these models to simulate the EAP waveform and, in such manner, create EAP-templates linked to ground-truth, specific electrophysiology-, morphology- and Cre-reporter-based cell classes.

We show simulations for two example single-cell models, one SP (**Fig. 3a**) and one AP (**Fig. 3b**). Somatic action potentials were evoked via simulated convergent, Poisson-style synaptic input along the dendritic arbor (**Fig. 3a-b**). The simulated EAP from the model exhibits its largest amplitude in the somatic region and actively propagates into the dendrites. As for extracellular recordings, one- and multi-channel features of AP and SP were calculated from the simulated EAP waveforms. We see that the trough-to-peak width (TPW) and repolarization time (REP) of the simulated cells are very similar to the ones from experimental recordings (**Fig. 3c**). Furthermore, cell class differences predicted by simulations agree with *in vivo* recorded EAPs, e.g. simulated AP cells exhibit significantly lower TPW (two-sample t test, p=0.00025) and REP (Mann-Whitney U test, p=0.00024) than SP ones (**Fig. 3d**). Furthermore, because the biophysical models agree with experimental recordings for one-channel features TPW and REP, they can be used to link between *in vitro* properties of cell class and *in vivo* EAPs. We asked whether the experimentally measured intrinsic properties of the actual cells each model represents differentiate between morphology class AP and SP. Comparison between *in vitro* cellular data used to develop each of the AP and SP models (same mouse IDs as **Fig. 3c-d**) show statistically significant differences in intrinsic properties known to differentiate between major excitatory and inhibitory classes (spike width, adaptation, spike rate and f-I slope; **Fig. 3e**). We conclude that, not only the models, but also the underlying *in vitro* experiments mapping on RS and FS clusters, exhibit robust separation in slice electrophysiology properties known to separate excitatory from inhibitory classes.

**Figure 3.**
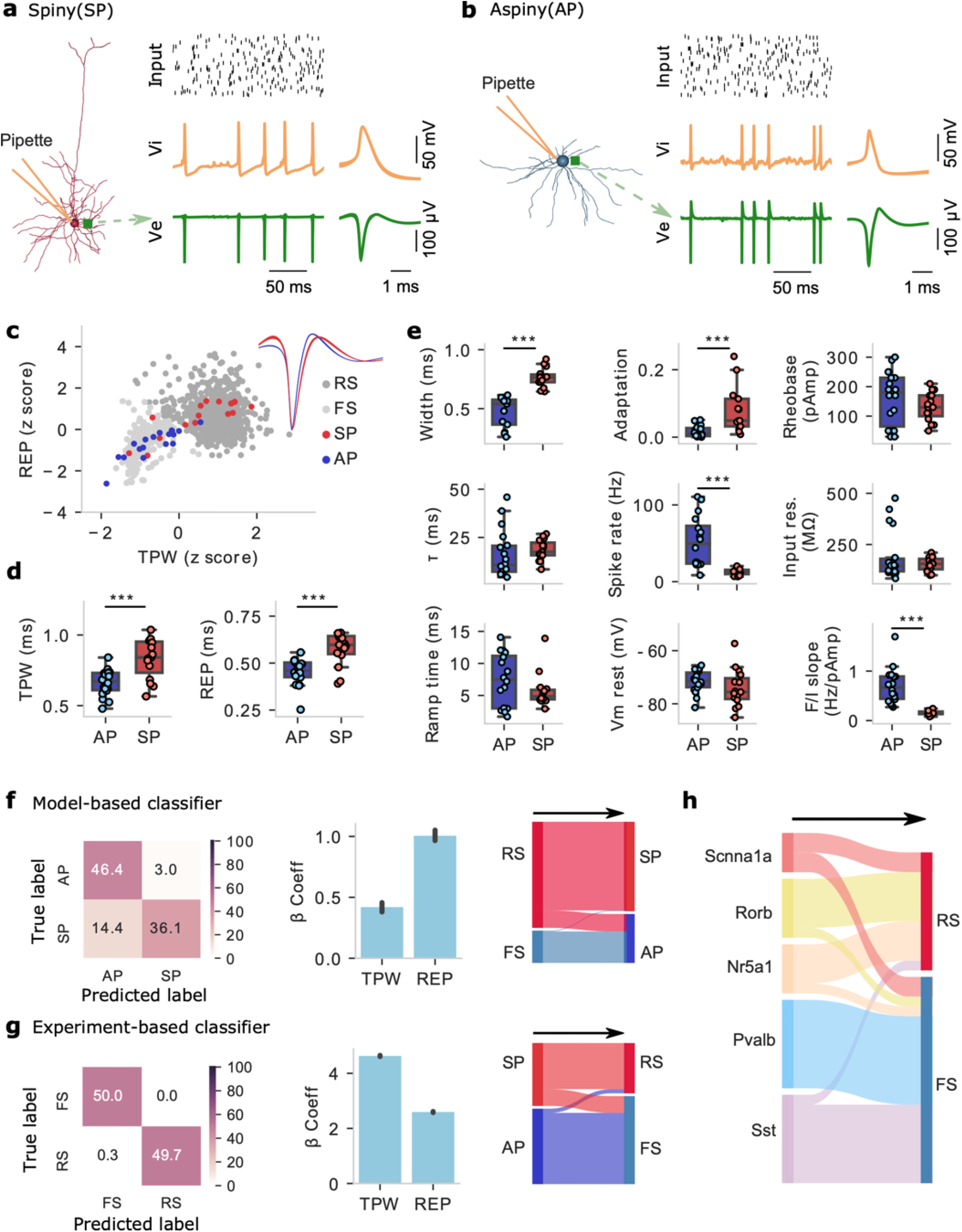
Classification of one-channel EAP (extracellular action potential) features of single-cell models and correspondence to *in vitro* data modalities. **a-b)** Bio-realistic single-cell models (one aspiny, AP, panel a; one aspiny, SP, panel **b)** activated via synaptic activity along their reconstructed dendrites result in spiking. Top: synaptic input (black bars: spike raster plot); Middle: intracellular voltage Vi trace (orange); Bottom: extracellular voltage Ve (green) close to the soma (location designated by the green square). Time traces (left) and mean Vi and EAP waveforms (right). **c)** One-channel EAP analysis from single-cell models (*n*=33, blue: AP; red: SP) and *in vivo* units (light grey: fast-spiking (FS) units; dark grey: regular-spiking (RS) units). **d)** Comparison of TPW (trough-peak width, two-sample t-test, two-sided, p=0.00025) and REP (repolarization time, Mann-Whitney U test, two-sided, p=0.00024) from simulated EAP waveforms between AP (n=18) and SP (n=15) models. Box plots show center line as median, box limits as upper (75%) and lower (25%) quartiles. The whiskers extend from the box limits by 1x the interquartile range. ***p<0.001. **e)** Comparison of intrinsic properties extracted from *in vitro* Vi dynamics between the AP (n=18) and SP (n=15) neurons (also used to generate the single-cell models). Mann-Whitney U test (two-sided) was used for width, adaptation, 𝜏, input res. (resistance), ramp time, and F/I slope; two-sample t-test (two-sided) used for rheobase, spike rate, and Vm rest (resting potential). ***p<0.001. **f)** Model-based classifier: classifier trained on one-channel EAP features (TPW, REP) of single-cell models to discriminate between AP (n=18) and SP (n=15) neurons (left: confusion matrix; middle: beta coefficients of the linear SVM classifier, bootstrap sampling 100 times; right: Sankey diagram showing the prediction on the test dataset). **g)** Same layout as in f, experiment-based classifier: classifier trained on one-channel EAP features (TPW, REP) of *in vivo* units to discriminate between FS (n=281) and RS (n=923) populations labeled via *K*-means clustering. **h**) One-channel EAP features of single-cell models (model labels: Cre-reporter lines, 4 Scnn1a, 6 Rorb, 5 Nr5a1, 9 Pvalb and 9 Sst) classified as FS or RS by using the experiment-based classifier. Source data are provided as a Source Data file.

To link between labels of *in vivo* units (RS vs. FS) and the morphology classes of simulated neurons (spiny or SP vs. aspiny or AP), we used a two-way classification process^27^. In one direction, the model-based classifier was trained on one-channel EAP features (TPW, REP) of models to discriminate between SP and AP neurons. This process yielded 82.5% classification accuracy on the validation data set (support vector machine, SVM; training/validation set, 75%/25%; **Fig. 3f**). Then, the model-based classifier was applied on the test data set (*in vivo* clustered FS and RS units from V1). Most FS units are labeled as AP neurons while the majority of RS as SP (**Fig. 3f**). We also tested the opposite direction. In the experiment-based classifier, we trained on one-channel EAP features (TPW, REP) of *in vivo* units to discriminate between FS and RS clusters (training/validation set, 75%/25%) and classification accuracy on the validation data set exceeded 99% (SVM; **Fig. 3g**). When applying the classifier on the test datasets, i.e., model-labeled AP and SP neurons, most AP neurons were labeled as FS and most SP neurons as RS units (**Fig. 3g**). We conclude that the majority of *in vivo* RS map to *in vitro* SP cells while the majority of *in vivo* FS map to *in vitro* AP cells based on one-channel features TPW and REP.

Beyond the intrinsic properties and morphology classes, the simulated neurons also contain Cre-line labels from the Cre-lines used *in vitro* to target the individual cells. In a subsequent analysis, instead of using the morphology labels SP and AP, we used the transgenic line label (excitatory: Scnn1a, Rorb, Nr5a1; inhibitory: Pvalb, Sst)^1^ of the models as input to the experiment-based classifier to predict the one-channel *in vivo* clusters (RS vs. FS). The excitatory classes (Scnn1a, Rorb and Nr5a1) are mainly classified as RS whereas inhibitory classes (Pvalb and Sst) are mainly classified as FS (**Fig. 3h**). We conclude that the biophysical models agree with experimental *in vivo* EAP recordings in terms of one-channel EAP features and reflect experimental intrinsic and morphology class-dependent differences also observed *in vitro*.

### Composition and properties of multi-channel RS clusters

We next attempt to deduce single-cell intrinsic electrophysiology and morphology properties of the *in vivo* multi-channel clusters. We first asked whether the single-cell models recapitulate the three multi-channel clusters for each class. Starting with the SP models, we clustered the models based on their simulated multi-channel EAP features (n=15 SP models; *K*-means clustering). Two separate clustering analyses (elbow method and the density function) determined the number of SP clusters in our simulated data to be three, i.e. SP1-3. Notably, the number of SP clusters coincides with the number of RS clusters detected *in vivo* (RS1-3) (**Fig. 4a**). Among three RS clusters, there is no significant difference in the largest amplitude channel (**Fig. 4a, right**), however, the waveform propagation separates them into three clusters (**Fig. 4b**). Looking at the multi-channel features (1/*V_below_* and 1/*V_above_*) there is correspondence between SP1 with RS1, SP2 with RS2 and SP3 with RS3. This is also reflected in the distinct EAP propagation properties of the three SP clusters, with SP1 showing faster supragranular propagation than SP2 while SP3 shows reduced infragranular propagation vs. SP1 (**Fig. 4b**). We conclude that the biophysical models of morphologically spiny neurons SP separate into three distinct clusters (SP1-3) based on the same multi-channel features that also separate *in vivo* multi-channel RS units into clusters RS1-3 with EAP propagation patterns that resemble model and *in vivo* clusters.

**Figure 4.**
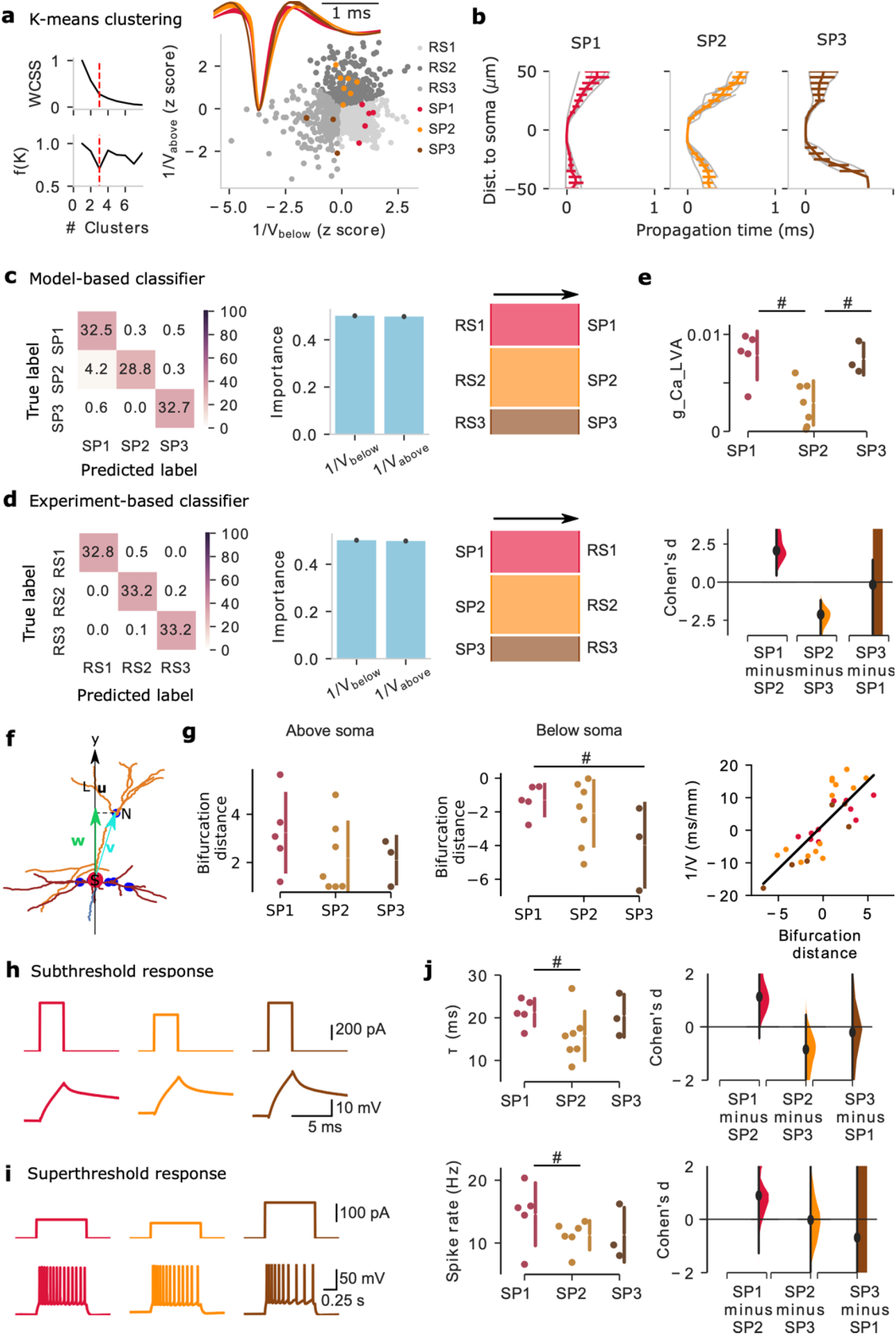
Distinct cellular properties of multi-channel regular-spiking (RS1-3) clusters. **a)** Clustering of spiny (SP) models using *K*-means clustering based on multi-channel extracellular action potential (EAP) features. Both the elbow method and density function analysis independently identify three multi-channel SP clusters (left: within cluster sum of squares (WCSS) and density function, broken red line: optimal number of clusters; right: model-based SP clusters; inset: mean EAP-waveform of each RS-population). SP1-3 and RS1-3 are shown using the multi-channel features 1/V_below_ and 1/V_above_ (the inverse of spike propagation velocity below/above soma location). **b**) Spike propagation along the simulated probe as function of distance from the soma (channel with largest EAP amplitude) for the three SP classes, SP1-3 (grey lines: propagation of individual models; n=5 SP1, n=7 SP2, n=3 SP3; colored lines: mean ± SD (standard deviation)). **c**) The model-based classifier (random forest) trained on the multi-channel features (1/V_below_ and 1/V_above_) identifies SP1-3 (left: confusion matrix; middle: feature importance based on classifier; right: sankey diagrams show the prediction on the test dataset). **d**) Same layout as in **c**, the experiment-based classifier was trained on multi-channel *in vivo* EAP features to discriminate between RS1-3. **e**) Comparison between model conductances ascribed to SP1-3. The largest effect size across the conductances is found for axonal Ca_LVA. # indicates Cohen’s d effect size >0.8. **f)** Bifurcation distance (**w**) of one bifurcation node in the reconstructed morphology of a neuron is defined as the projection of the vector (**v**) from soma (S, red dot) to the position of the bifurcation node (N, blue dot) projected to a line (**u**) connecting the soma (S) to a node (L) in y axis. **g**) Morphology bifurcation distance above soma (left) and below soma (middle). Right: inverse of wave propagation velocity vs. the bifurcation distance (line: linear fit). **h-i**) Intrinsic properties from *in vitro* experiments based on SP1-3 (subthreshold and spiking responses). **j**) Comparison of cellular time constant (𝜏) and max spike rate (response to dc current injections) among *in vitro* experiments based on SP1-3. # indicates Cohen’s d effect size >0.8. Source data are provided as a Source Data file.

We looked deeper into the correspondence between the model-based SP1-3 and *in vivo* clusters RS1-3 defined via the multi-channel EAP features by using two-way classification: supervised classifiers trained on the simulated EAPs of modeled neurons then applied to *in vivo* units (“model-based classifier”), and supervised classifiers trained on experimental *in vivo* units, then applied to the model classes (“experiment-based classifier”). Specifically, the model-based classifier trained on multi-channel EAP features (1/*V_below_* and 1/*V_above_*) to identify SP1-3 showed excellent performance (random forest; classification performance >94%; **Fig. 4c**). In a next step, we applied the model-based classifier on the test experimental data sets (*in vivo* clustered RS1-3) and found that, indeed, RS1 units are mapped to SP1, RS2 to SP2, and RS3 to SP3 with high fidelity (performance: >94%; **Fig. 4c**). We also pursued the opposite direction by building the experiment-based classifier trained on multi-channel *in vivo* EAP features to discriminate among RS1-3 and saw very high classification accuracy (> 99%; **Fig. 4d**). The experiment-based classifier on the test simulation data sets (models clustered SP1-3), once more, cleanly maps SP1 to RS1, SP2 to RS2 and SP3 to RS3, respectively (**Fig. 4d**). Thus, our initial results are validated by the two-way classification that robustly maps model-based SP1-3 classes to *in vivo* RS1-3 clusters via their multi-channel features.

Since RS1-3 are mapped to SP1-3, respectively, what other properties of the *in vivo* clusters RS1-3 can be deduced from the SP1-3 data and associated models? We address this question for three data modalities: models, morphologies and intrinsic electrophysiology properties. First, we asked whether SP1-3 models can point to key differences between the three clusters in terms of the conductance setup. Pairwise comparison between SP1-3 model conductances indicates that the axonal low-voltage activated Ca-conductance is increased for SP1 and SP3 vs. SP2 (Cohen’s d effect size > 0.8; **Fig. 4e**), i.e. a conductance linked to elevated spike rate (bursting) and rapid spike recovery^37^. In terms of cellular morphology, given SP1-3 have different spike propagation profiles, we used a morphology feature looking at the cable structure attached to the soma, the bifurcation distance. The bifurcation distance is the normalized distance between the soma and the dendritic bifurcation with a large bifurcation distance effectively translating to a longer unobstructed path along the dendrite (see also Methods). Pairwise comparison of the bifurcation distance above soma and below soma among SP1-3 (the reconstructed morphologies were also used to develop the models) reveals differences in one property, the basal dendrite bifurcation distance below the soma (**Fig. 4f-g**; see also Methods). Specifically, SP1 and SP3 have different bifurcation distance especially below the soma (**Fig. 4g**). Notably, the morphology bifurcation distance, exhibits a strong linear relationship with the spike propagation speed across SP1-3 (**Fig. 4g** right, slope = 2.7, the correlation coefficient r = 0.8, p = 1.02*10^-7^). A larger bifurcation distance, results in a lower spike propagation speed along the basal (negative bifurcation distance) and apical (positive bifurcation distance) arbor. Thus, class-dependent morphology properties that impact spike propagation can also lead the class-dependent propagation speed and symmetry differences observed between SP1-3 (Fig. 5b). Finally, we compared *in vitro* subthreshold (**Fig. 4h**) and spiking (**Fig. 4i**) intrinsic electrophysiology properties among SP1-3 (the slice experiments also used to develop the models) and found differences in the cellular time constant 𝜏 and peak spike rate (response to dc current injections, **Fig. 4j**). Specifically, SP1 neurons achieve a higher spike rate especially compared to SP2, which, in turn, agrees with the model-based observation of increased axonal low-voltage activated Ca-conductance of SP1 (**Fig. 4e**). Moreover, SP1 is more electrotonically compact than SP2 (**Fig. 4j**). We conclude that, by virtue of mapping SP1-3 to RS1-3, the multimodal comparison between models (including their associated *in vitro* experiments) and *in vivo* clusters yields several distinct properties: a difference in axonal low-voltage activated Ca-conductance (SP1 and SP3 vs. SP2), a morphology difference in the basal dendrite bifurcation distance below the soma (mainly in SP1 vs. SP3) that, in turn, impacts the spike propagation speed, and, finally, SP1 being more electrotonically compact than SP2.

**Figure 5.**
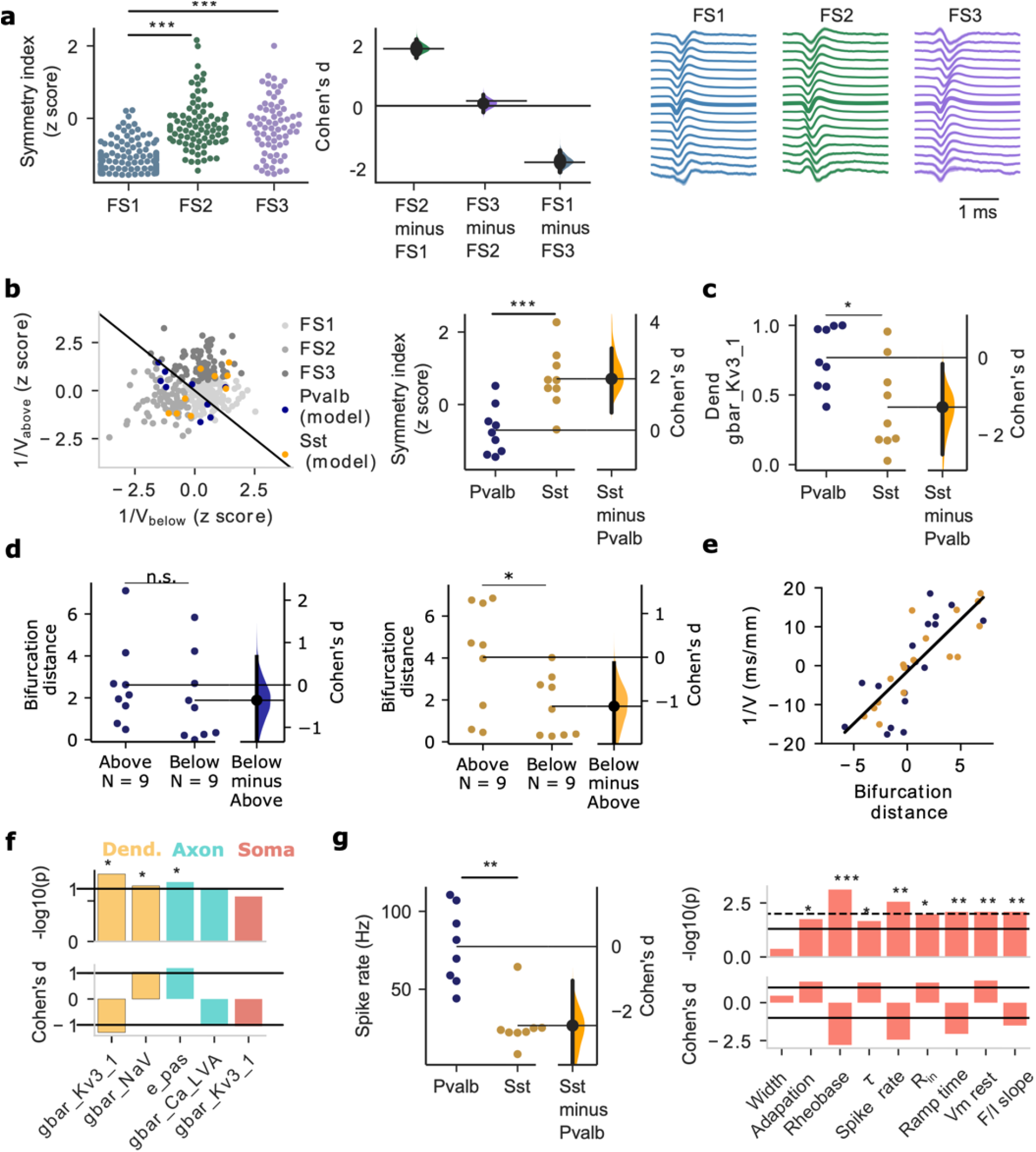
Distinct cellular properties of multi-channel FS clusters. **a)** The spike propagation symmetry index separates FS1 from FS2-3 (left, circles: experimental measurements; middle: effect size measured by Cohen’s d; right: mean spatiotemporal spike propagation of multi-channel clusters FS1-3; *n*= 130 FS1, *n*=82 FS2, *n*=69 FS3). Kruskal-Wallis H-test, F=111.41, p-values corrected using the Holm-Bonferroni method for multiple tests, ***p<0.001. Error bars represents a bootstrap 95% confidence interval. **b**) Left: the multi-channel features (V_below_, V_above_) clustering FS units (grey) and superposed multi-channel EAP features of models of Pvalb (blue, n=9) and Sst (yellow, n=9) neurons. Right: spike propagation symmetry index for Pvalb and Sst single-cell models. two-sample t-test, two-sided, p=0.000998. **c**) Pairwise comparison of model conductances between Pvalb and Sst models. The strongest and most statistically significant difference is shown in the dendritic Kv3.1 conductance. Mann-Whitney U test, two-sided, *p<0.05. **d**) Morphology bifurcation distance above (two-sample t-test, two-sided, p=0.45) and below (two-sample t-test, two-sides, p=0.03) soma between Pvalb (left, dark blue) and Sst (right, orange) models. *p<0.05. **e**) The inverse of spike propagation velocity vs. the bifurcation distance (line: linear fit; + indicates above soma, - indicates below soma). **f**) Pairwise comparison of Pvalb vs. Sst model conductances (top panel, -log10(p-value), black line: p=0.05; bottom panel, Cohen’s d effect size, black lines: |d|=0.8). The comparison of dendritic Kv3.1 conductance as shown in **c**. Mann-Whitney U test, two-sided, *p<0.05. **g**) Left: pairwise comparison between intrinsic properties of Pvalb and Sst neurons measured *in vitro* (same experiments as the ones used to develop to single-cell models). Maximum spike rate to dc current injections separates between Pvalb and Sst neurons. Mann-Whitney U test, two-sided, p=0.0067; Right: pairwise comparison between nine intrinsic properties of Pvalb vs. Sst neurons (top: statistical significance expressed in terms of -log10(p-value); solid line: *p*-value=0.05, broken line: p-value=0.01; bottom: Cohen’s d effect size, solid black line: |d|=0.8) also used to generate the computational models. Mann-Whitney U test, two-sided, *p<0.05, **p<0.01, ***p<0.001. Source data are provided as a Source Data file.

### Multi-channel features separate inhibitory Pvalb and Sst

FS units are most typically associated with inhibitory cell classes that are inherently heterogeneous. For example, Pvalb includes fast-spiking basket cells as well as Chandelier cells, while Sst includes Martinotti and non-Martinotti cells. We also found that this diversity of interneurons is reflected in FS1-3. While we focused our analysis on the two most populous inhibitory classes, Pvalb and Sst ^1^, we saw no clear mapping between FS1-3 and Pvalb/Sst. We therefore decided to introduce an additional multi-channel feature, the symmetry index (see Methods), quantifying the spatial characteristics of spike propagation and, in this manner, account for another aspect of morphology and its impact on the spike signature. Using the symmetry index to look at FS1-3 we saw a separation between FS1 (symmetric spike propagation) and FS2/FS3 (asymmetric spike propagation) (**Fig. 5a-b**). Notably, clearer separation between Pvalb and Sst models was achieved based on the symmetry index (**Fig. 5b**, right; n=9 Pvalb models, n=9 Sst models). We conclude that while multi-channel features 1/*V_below_* and 1/*V_above_* do not exhibit clear mapping, accounting for an additional multi-channel feature, the symmetry index, separates biophysical models of Pvalb and Sst.

Which properties can be deduced from the models and associated *in vitro* data? Once more we consider three data modalities: models, morphologies and intrinsic properties from the *in vitro* Pvalb (n = 9) and Sst (n = 9) experiments (**Table S1**). Pairwise comparison between Pvalb and Sst models at the level of ionic conductances reveals statistically significant differences in three conductances with the effect size being largest for Kv3.1 (**Fig. 5c, f**). Elevated Kv3.1 expression is a key differentiator between Pvalb and other inhibitory cell types, i.e. increased Kv3.1 results in a shorter spike width and fast afterhyperpolarization^24, 39–41^. In terms of cellular morphology, pairwise comparison of morphology features (bifurcation distance above and below soma) from the reconstructions in the Pvalb (**Fig. 5d**, left, dark blue) and Sst (**Fig. 5d**, middle, orange) cells show a statistically significant difference in the bifurcation distance between above and below soma in Sst cells. Specifically, while Pvalb morphologies are symmetric (i.e., above vs. below bifurcation distance remains similar), Sst possess a more asymmetric morphology with the bifurcation distance above being longer than below their soma (**Fig. 5d**). To look at how the bifurcation distance affects spike propagation, we plotted the bifurcation distance above (positive values) and below (negative values) against the spike propagation speed V in the model data. We found that the bifurcation distance above and below the soma is robustly related with the inverse of the EAP propagation velocity (**Fig. 5e**, right, slope = 2.67, the correlation coefficient r=0.8, *p*-value=5.12*10^-9^). Once more, a larger bifurcation distance results in a lower spike propagation speed. Thus, the increased bifurcation distance asymmetry leads to more asymmetric spike propagation along Sst morphologies. On the hand, the symmetry of Pvalb morphologies with respect to bifurcation distance leads to more symmetric spike propagation. Pairwise comparison of *in vitro* intrinsic electrophysiology properties between Pvalb and Sst (from the same experiments used to develop the Pvalb and Sst experiments) reveals several differences in peak spike rate, rheobase, resting potential (**Fig. 5g**) among others supporting that Pvalb are more electrotonically compact compared to Sst, which agrees with the observation about differences in Kv3.1 difference (**Fig. 5c, f**). In summary, the comparison between Pvalb and Sst models, morphologies and intrinsic properties points to a difference in Kv3.1, in bifurcation distance and in a several intrinsic properties shown to separate between Pvalb vs. Sst (e.g., peak spike rate) and shape intracellular dynamics as well as the EAP waveform.

We also examined whether differences in spike propagation symmetry between Pvalb and Sst can be attributed to morphology orientation. While the elongated somadendritic axis of pyramidal neurons can give rise to spike propagation asymmetry^42^, the impact of the angle between an extracellular probe and the cellular morphology of inhibitory cells remains unknown. In a separate series of simulations, we varied the angle between the extracellular probe and morphology across Pvalb and Sst models and found that, indeed, certain EAP multi-channel metrics including the symmetry index are affected by this parameter with certain constellations exacerbating the pairwise difference between Pvalb and Sst (**Fig. S8**). Even so, the robust and highly significant differences in symmetry index found between inhibitory classes can hardly be a mere reflection of rotation effects. While we cannot exclude this parameter contributing to the trends observed, the evidence clearly points to biophysical differences between the clusters rather than aspects of experimental layout. We conclude that Pvalb are distinct from Sst across multiple *in vitro* modalities considered in our work, a fact also reflected in their distinct EAP signatures that allows their *in vivo* identification and separation using multi-channel EAP properties.

### Comparisons with ground-truth channelrhodhopsin-tagged Pvalb and Sst units *in vivo*

So far, we deduced cellular properties of *in vivo* units by comparing the simulated EAP waveform from models linked to specific *in vitro* experiments of known identity to *in vivo* recorded EAP waveforms, and vice versa. Opto-tagging is a method that can link EAP measurements to specific cell types by directly photo-stimulating cells that express the light-activated channel channelrhodopsin-2 (ChR2) to a restricted neuronal subpopulation under genetic control^43, 44^. Opto-tagging experiments can thus offer ground-truth data with recorded EAPs originating from known populations of neurons. Here, we used a channelrhodopsin reporter line (Ai32) crossed with a driver line in which Cre recombinase expression was driven by Pvalb or Sst promoter (**Fig. 6a**, dark green region). This process resulted in ChR2-tagged Pvalb and Sst neurons that responded to light stimulation with short latency and reliably (**Fig. 6b**). Extracellular recordings with Neuropixels in these animals detected 25 well-isolated Pvalb units in 8 Pvalb-Cre mice and 18 Sst units in 12 Sst-Cre mice (**Fig. 6c**; see Methods; Supplementary Data 3).

**Figure 6.**
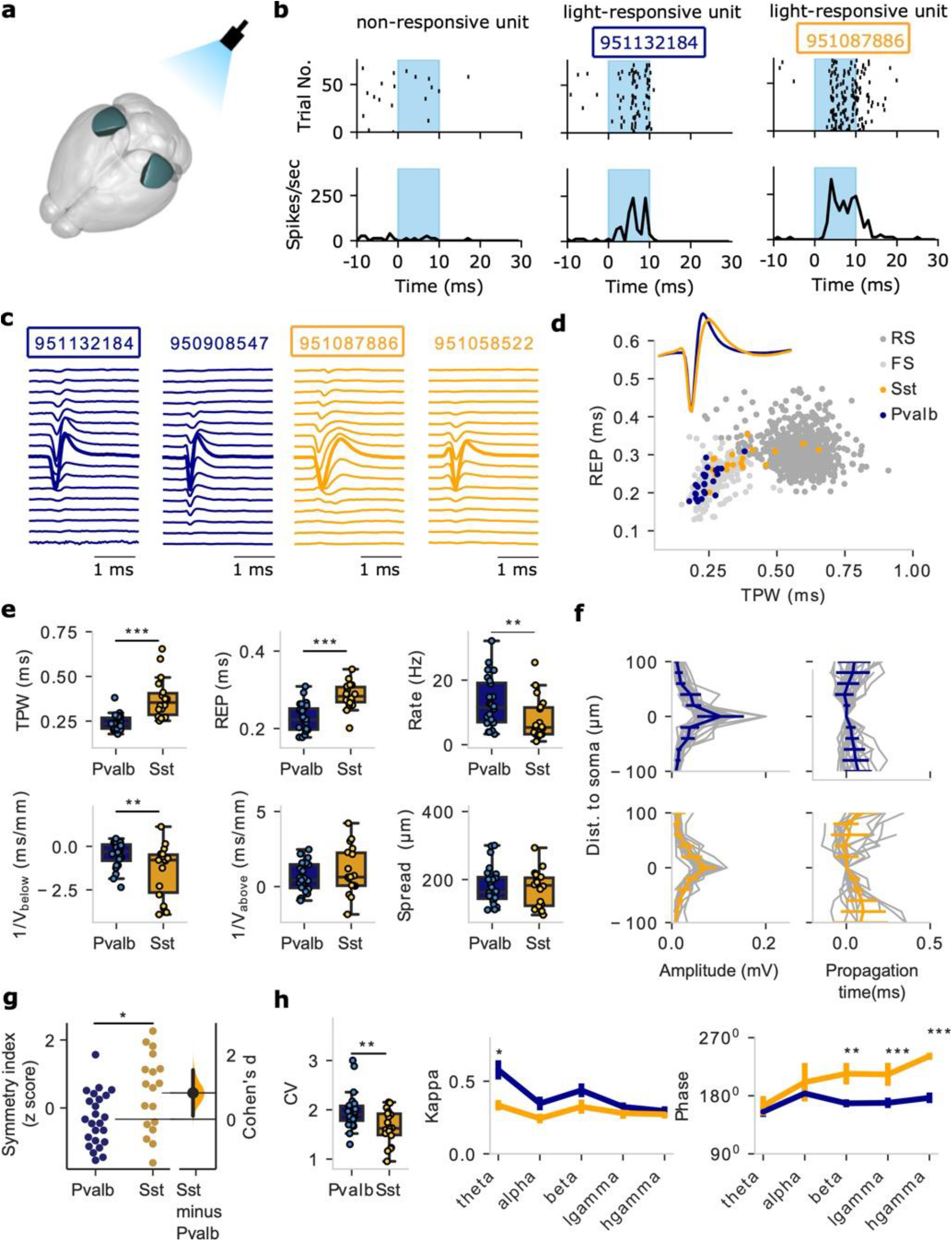
*In vivo* extracellular action potential (EAP) and functional properties of opto-tagged Pvalb and Sst neurons. **a)** Light sensitive channelrhodopsin-2 (ChR2) channels were virally expressed in two inhibitory cell populations, Pvalb and Sst, in mouse V1 (dark green areas). The animals were then implanted with Neuropixels probes. **b)** Example units responding to light activation (light blue regions) in V1. Top: spike rasters; Bottom: spike frequency. Left: a non-responsive unit; Middle: a light-responsive Pvalb unit; Right: a light-responsive Sst unit. **c**) Examples of multi-channel EAPs of Pvalb units (dark blue) and Sst units (orange). Two of the units are the same ones as in panel **b** (boxes). **d**) One-channel EAP features (trough-peak width: TPW; repolarization time: REP) for the Pvalb (dark blue, n=24) and Sst units (orange, n=18) from the optotagging experiments (inset: mean EAP waveforms; light gray: FS units, dark gray: RS units, from wild-type animals as in Fig. 2d). **e)** Comparison of EAP properties between optotagged Pvalb (n=24) and Sst (n=18) units (top: one-channel properties; bottom: multi-channel properties). Box plots show center line as median, box limits as upper (75%) and lower (25%) quartiles. The whiskers extend from the box limits by 1x the interquartile range. Mann-Whitney U test, two-sided, **p<0.01, ***p<0.001. **f**) EAP amplitude (left) and propagation (right) along the extracellular channels as function of distance from the soma (taken as the channel with the largest EAP amplitude) for the optotagged Pvalb (n=24) and Sst (n=18) units (gray lines: individual units; colored lines: mean ±SD (standard deviation)). **g**) Comparison of the symmetry index for Pvalb (n=24) vs. Sst (n=18) units (two-sample t-test, two-sided, p=0.012). **h)** Left: Comparison of response pattern during drifting gratings in the opto-tagging experiments (CV: coefficient of variation). Box plot representation is similar as in panel (**e**). Mann-Whitney U test, two-sided, p=0.0076, n=24 Pvalb, n=18 Sst; Right: spike-field coherency metric kappa and preferred spike phase of optotagged Pvalb and Sst for various LFP frequency bands. Data are presented as mean ± SEM (standard error of mean). Mann-Whitney U test, two-sided, *p<0.05, **p<0.01, ***p<0.001. Source data are provided as a Source Data file.

Using this ground-truth data set for two major inhibitory cell classes we pursued one- and multi-channel EAP analysis. For one-channel EAP features (TPW, REP), the opto-tagged Pvalb units exhibit clear overlap with FS from experiments with wild-type animals. Sst units are much more diffuse spanning across the FS/RS-space (**Fig. 6d**). Direct comparison of one-channel features (TPW, REP) and *in vivo* activity metrics like spike frequency show that Pvalb are well-separated from Sst (**Fig. 6e**, top). Pvalb and Sst also exhibit clear differences in terms of multi-channel EAP propagation, especially when looking at the symmetry index. Specifically, the optotagged recordings reveal that Pvalb exhibit symmetric and fast propagation profile while Sst exhibit less symmetric propagation and increased variability (**Fig. 6e-f**). Pairwise comparison of the symmetry index for the Pvalb and Sst optotagged units confirms that Pvalb show more symmetric EAP propagation compared to Sst (**Fig. 6g**), in agreement with simulations (**Fig. 5b**). We conclude that *in vivo* ground-truth opto-tagging experiments show that Pvalb and Sst are separable in terms of one- and multi-channel properties (symmetry index) in line with findings from the computational models.

We also looked for functional differences between Pvalb and Sst in the opto-tagged units. First, we found that Pvalb exhibit higher spike time variability than Sst (**Fig. 6h**, left). More interesting differences appear for phase-locking to ongoing LFP oscillations. Specifically, we found that Pvalb exhibits stronger phase-coupling than Sst for slower (theta) oscillations. Furthermore, Pvalb have a significantly different spike phase especially for faster oscillations (beta, low- and high-gamma) than Sst with Sst units spiking in a later phase by about 40-50°. (**Fig. 6h**). We note the similarity of this pattern with the spike phase relationship of wild-type units FS1 and FS2 (**Fig. 2e**). We conclude that the opto-tagging experiments reveal that, beyond separable in terms of multi-channel features, Pvalb units also have more variable spiking as well as stronger coupling to theta and earlier spiking for faster oscillations compared to Sst.

## Discussion

Understanding the role and function of cellular taxonomies in behavior is an important challenge in an era where advancements in sequencing technologies continuously refine these taxonomies^1–4, 9, 10^. Extracellular electrophysiology recordings offer unparalleled ability to monitor cellular activity *in vivo* across spatiotemporal levels yet lack cell type-specificity, with optotagging making it possible to label only one or two distinct cell types per experiment^21, 45, 46^. Here we introduce a framework for the identification and characterization of major cortical cell types solely based on their extracellular electrophysiology signatures with multiple data modalities. Our starting point are EAP waveforms recorded from high-density Neuropixels probes in mouse primary visual cortex (V1). Using one-channel EAP features we separated units into two clusters, FS and RS, that exhibit differences both in terms of EAP waveform and functional properties such as LFP entrainment. We separately looked at phase coupling in prominent LFP oscillation bands (theta, alpha, beta, low and high gamma) and found that FS units are consistently more entrained across LFP-bands compared to RS units. In agreement with other studies (e.g. ^13^), FS and RS exhibit significantly higher phase-locking during drifting gratings than during spontaneous activity across LFP-bands. When we looked at the preferred spike phase, we found that FS spiking came earlier in the cycle than RS in beta and low gamma. These observations are in line with the studies of neocortical unit activity in humans and monkeys^35^. Specifically, FS phase precedence is also in line with^35^ and opposite to the hippocampal activation pattern observed during high frequency ripples^36^. When we looked at phase coupling along the cortical depth, we found a diverse landscape. While FS remain consistently more entrained than RS, FS phase precedence over RS is spatially inhomogeneous and particularly pronounced in the granular region (broadly layer 4) for beta, low and high gamma. In contrast, in the supragranular and infragranular regions, FS and RS clusters exhibit less pronounced phase differences across LFP-bands despite significant differences in coupling strength. We conclude that the FS and RS clusters represent larger families of diverse cell classes organized along the cortical network serving different roles *in vivo*.

Expanding the feature set from one-channel to multi-channel EAP features results in further separation within the RS and FS groups into six finer groupings, three FS (FS1, FS2, FS3) and three RS (RS1, RS2, RS3) clusters. We show that the six clusters exhibit functional differences in their dynamics to visual stimuli (e.g., drifting gratings in head-fixed animals) and differential coupling to ongoing LFP oscillations. Looking at the properties of these finer clusters with cortical depth we found increased diversity in their spike-LFP coupling. RS3, for example, exhibits almost double the coupling strength than RS1 in the supragranular region for beta and low gamma (RS2 is an intermediate case). On the other hand, in the infragranular region and for low gamma, RS3 and RS1 spike phase is similar while RS2 comes earlier (the same happens for high gamma). The differences in RS1-3 are consistent with classes of neurons that possess different biophysical setups as well as a divergence in connectivity patterns. It is known, for example, that the biophysical properties of excitatory V1 neurons vary and depend on cortical depth which, in turn, is expected to have an impact on their firing properties and burstiness ^34, 47–49^. In addition, their intricate connectivity and projections along the anatomical hierarchy can result in a spectrum of functional clusters among excitatory V1 cells that reflect upstream input segregation from earlier brain regions (e.g., various thalamic areas ^50–52^). The combination of diverging biophysical properties of V1 excitatory cells combined with localized and class-specific connectivity gives rise to functionally distinct and input-specific RS1-3 clusters. Furthermore, behavior and brain state can further modulate the response properties of excitatory clusters along V1^50^. The aforementioned points to a network consisting of clusters of distinct biophysical properties and functional *in vivo* responses that, nevertheless, can be organized and reconfigured in multiple ways, depending on the external input and internal state.

While excitatory cells exhibit differences in visual responses (though with varying degree of sensitivity and selectivity), inhibitory neurons do not show strong or selective responses confirming observations using the same visual inputs^53^. Even so, they play a central role in shaping cortical activity in terms of orchestrating and patterning ongoing and/or evoked oscillations^7, 38, 54–56^. Indeed, when we looked at the phase-coupling properties of FS1-3 we found differences in the alpha, beta, and gamma bands. Furthermore, in an additional analysis we observed differences between FS1-3 (mainly in the gamma bands) as function of cortical depth. While the diversity of inhibitory coupling to ongoing oscillations remains elusive in V1 (though see^33, 50^). In agreement with other V1 studies^50, 57^, our experiments support the observation that the most prominent LFP pattern in the waking V1 is a theta-band oscillation (hypothesized as an evolutionary precursor of the primate alpha activity in the visual cortex). Yet, we also found that FS1-3 (but also RS1-3 as well as FS-RS) differentiate their coupling in higher LFP-bands, i.e., in alpha, beta and gamma, rather than in the band of their most prominent pattern (theta). Notably, inhibitory parvalbumin- and somatostatin-positive interneurons exhibit large amplitude, rhythmic hyperpolarization at 3–6 Hz in V1 during behavior^50, 57^.

The distinct properties of FS1-3 in EAP waveform and coupling strength/phase to LFP oscillations is reminiscent of the distinct hippocampal inhibitory classes and their coupling to local theta, gamma and sharp wave ripples, e.g.^36, 55, 56, 58–60^. For example, two putative inhibitory classes located in the pyramidal layer and the alveus/stratum oriens of hippocampal CA1 with distinct EAP waveforms also exhibit differences in discharge probability and theta spike phase with one coming earlier by about 30^0^ and both preceding pyramidal spiking^36^. The picture is reversed during ripple activity when phase differences between the two inhibitory clusters are minimized and pyramidal spiking precedes both^36^. *In vivo* recordings combined with tedious morphological characterization unravel distinct coupling features, e.g. between palvalbumin-expressing basket cells, bistratified and cholecystokinin-expressing interneurons differing their spike phase by 30^0^-40^0^ during the gamma cycle^60^. The coupling strength as well as the phase differences observed between distinct cell classes during oscillations are in line with what we see for FS1-3. Excitatory pyramidal neurons in hippocampus and neocortex also form distinct morphological, molecular, connectivity and functional populations ^61–64, 64–68^. A major organizing principle of excitatory neurons is cortical depth and the presence of functionally distinct sublayers ^69^ – in CA1, this organization is also reflected in the cellular and functional properties with deep cells spiking faster, burstier and exhibiting stronger modulation for slow oscillations ^70, 71^. Neocortical organization is less understood with respect to its functional modules and their role in oscillations (although see ^72–76^) yet the RS1-3 coupling profile points to the existence of a cellular and functional organization along the depth axis.

To map between the cellular *in vitro* classes and subclasses and *in vivo*, EAP-based clusters, we develop biophysical models that reflect key properties of *in vitro* cell types and use these models to simulate EAP properties. We use a computational optimization workflow to generate and evaluate biophysically realistic, cell type-specific cellular models with active conductances at scale^24^. We then use two-way classification to map *in vitro* classes to *in vivo* clusters and vice versa, with models providing the link between the two worlds and the associated class/cluster label. In a stepwise manner, we show that a set of one-channel EAP features (TPW, REP) separates *in vivo* EAP clusters in terms of spike rate (FS vs. RS units) and *in vitro* morphology classes (AP vs. SP neurons). The fact that narrow EAP waveform units map to FS and AP while wide units map to RS and SP is in line with previous work^14–18, 27^. A fraction of simulated excitatory neurons also mapped onto FS units that we attribute to some excitatory classes that possess narrow spike width and some model discrepancy that prohibits capturing all EAP features in their full extent. The latter can lead, in a few cases, to mislabeling. Yet, it is the use of these models that also enables linking seemingly disparate data sets in a manner that results in specific and testable hypotheses about the identity and properties of the various clusters (e.g. in terms of the underlying conductance or morphology differences between the *in vivo* clusters).

Looking at RS1-3, we found that RS1/SP1 and RS3/SP3 are electrotonically more compact than RS2/SP2 with a possible biophysical mechanism accounting for such differences being the axonal low-voltage activated Ca-conductance. Moreover, we found that basal dendrite differences between RS2/SP2 and RS3/SP3, a feature that could potentially explain the EAP waveform symmetry between RS1-3 clusters. For FS1-3, we found that biophysical models of Pvalb and Sst broadly capture the multi-channel properties of FS1-3 and specifically the distinct symmetry of FS1 vs. FS2-3 spike propagation. Notably, Sst cells are diverse in their morphology, which resulted in a wider range of multi-channel features. Comparing Pvalb and Sst models, morphologies and intrinsic properties we found a difference in Kv3.1, in bifurcation distance and in a several intrinsic properties shown to separate between Pvalb vs. Sst (e.g., peak spike rate) and shape intracellular dynamics as well as the EAP waveform. A set of *in vivo* ground-truth opto-tagging experiments validated that Pvalb and Sst are separable in terms of one- and multi-channel properties further supporting our observations based on computational models.

Our study shows that multi-channel EAP features can critically contribute to the separation of meaningful *in vivo* clusters. The key data modality reflected in these multi-channel properties is the cellular morphology^22, 25, 26^. It follows that for computational models to account for such properties, they need to account either for the fully reconstructed morphology^24^ or, at the very least, for key aspects of it^42^. Moreover, ionic mechanisms along the dendritic morphology also impact spike propagation intracellularly^77, 78^ and extracellularly^25, 26, 32^ pointing to an interesting possibility: the use of optotagging experiments to measure cell type-specific (e.g. Pvalb and Sst) multi-channel EAP properties *in vivo* and, in a second step, using these properties to constrain model parameters along the dendritic arbor where intracellular data is challenging to collect.

Notably, while Neuropixels recordings result in large numbers of recorded units, the bottom-up approach (i.e., generating data from transgenic lines *in vitro* by whole-cell patch-clamp and morphology reconstructions of labelled neurons) is a lower-yield and labor-intense process. In addition, the computational framework to turn the *in vitro* data (features of electrophysiology traces in combination with reconstructed morphologies) into biophysically realistic all-active single-cell models involves computationally intensive multi-objective optimization procedures (see Methods). This results in a natural imbalance in our data sets: a large number of isolated *in vivo* units compared to a smaller number of *in vitro* recorded and reconstructed neurons and models. Ever-increasing availability of high-quality, annotated cellular electrophysiology, morphology, transcriptomics data – the precondition to generate faithful, cell type-specific computational models at any scale – is underway and is expected to tackle the imbalance between the number of cellular models and *in vivo* recorded units. The larger the number of models and cell classes reflected in them, the better and more refined classifiers can be trained to map *in vitro* types to *in vivo* EAP clusters. With increasing cellular data and single-cell model availability, increasingly finer classification of EAP signatures can be achieved across different brain areas and, even, species that allows deducing cellular and functional differences between cell classes across data modalities.

## Methods

### *In vivo* Neuropixels recordings

All *in vivo* recordings come from the Allen Brain Observatory Visual Coding Neuropixels dataset^23^, accessible via the AllenSDK (https://allensdk.readthedocs.io/en/latest/visual_coding_neuropixels.html) and the DANDI Archive (https://gui.dandiarchive.org/#/dandiset/000021). Recordings were performed in awake, head-fixed mice allowed to run freely on a rotating disk. During the recording, mice either passively viewed visual stimuli (flashes) or viewed a mean-luminance gray screen. Data were collected from 25 wild-type C57BJ/6J mice (24 male, 1 female), and 8 Pvalb-IRES-Cre (6 male, 2 female) and 12 Sst-IRES-Cre (8 male, 4 female) crossed with an Ai32 channelrhodopsin reporter line^79^. Cre+ cells from Ai32 lines are highly photosensitive, due to the expression of Channelrhodopsin-2^80^. The Neuropixels probe can record from 384 contacts across 3.84 mm of tissue coverage (selectable from 960 available sites on a 10 mm length shank). In this study, we analyzed recordings from the primary visual cortex (V1). All extracellular spike data were acquired with Neuropixels probes^21^, with 30 kHz sampling rate (which achieves 0.033 ms temporal resolution) and a 500 Hz analog high-pass filter. Spike times and waveforms were automatically extracted from the raw data using KiloSort2^81^.

### Biophysical realistic all-active single-cell models

We use the biophysically realistic all-active single-cell model for 18 aspiny (AP) and 15 spiny (SP) mice neurons. The all-active models contain active conductances along the entire neuronal morphology. The dendritic arbors are adopted in the models from the reconstructed morphology. The models were generated with a computational optimization pipeline (**Fig. S2**) aiming for models that reproduce the intrinsic firing patterns and spike properties of individual neurons from two data modalities: the reconstructed morphology and the somatic electrophysiology response from *in vitro* whole-cell patch-clamp experiments. The models were fit with several voltage-gated sodium, potassium, and calcium conductances expressed at the cell soma, axon, and dendrites, using data from individual neurons in the Allen Cell Types Database (http://celltypes.brain-map.org/data). The optimization pipeline (multi-objective genetic optimization) was used to optimize the conductance densities by training the models with experimental somatic recordings in response to step currents^24^. The active conductances and passive properties marked according to their inclusion in each of the morphology sections (apical, basal dendrites, soma and axon) are reported in **Table 1**. We optimized both the spiking properties of the cell model (spiking timing, spike rate, etc.) given a particular morphology and features of the intracellular action potential waveform (spike amplitude, width, etc.) Only the models that passed certain criteria (Tol = 0.5 for both spike amplitude and spike width) were selected, where Tol is the tolerance. Specifically, the spike amplitude of the model should be in the range of [1-Tol, 1+Tol]*A_exp, while the spike width of the model should be in the range of [1-Tol, 1+Tol]*W_exp, where A_exp, W_exp represent the spike amplitude and width from experiments.

**Table 1.** Inclusion of each parameter in the morphology sections

| Parameters | Ra | $g_{pas}$ | $e_{pas}$ | $C_m$ | $I_h$ | Nav | $K_T$ | Kd | $K_v$<br>3.1 | $K_v$<br>2 | $I_m$ | SK | Ca<br>HVA | Ca<br>LVA |
| --- | --- | --- | --- | --- | --- | --- | --- | --- | --- | --- | --- | --- | --- | --- |
| Apical | √ | √ | √ | √ | √ | √ | × | × | √ | × | √ | × | × | × |
| Basal | √ | √ | √ | √ | √ | √ | × | × | √ | × | √ | × | × | × |
| Soma | √ | √ | √ | √ | √ | √ | × | × | √ | × | × | × | √ | √ |
| Axon | √ | √ | √ | √ | × | √ | √ | √ | √ | √ | × | √ | √ | √ |

**Table 2.** Stimulus metrics

| Stimulus | Metric | Description |
| --- | --- | --- |
| drifting gratings | modulation index | The phase-dependent responses to drifting gratings |
| | $f_1/f_0$ | The ratio of the 1st harmonic to the 0<sup>th</sup> harmonic |
|  | lifetime sparseness | The sparseness of individual neurons |

After a single-cell model is optimized, we simulated the extracellular potential using NEURON 7.5 simulator (https://www.neuron.yale.edu/neuron/) in combination with the Brain Modeling Toolkit (https://github.com/AllenInstitute/bmtk). This toolkit can simulate a variety of intracellular dynamics (e.g., spikes, and membrane voltages), as well as computing additional data modalities such as the extracellular potential. The extracellular potentials were computed using the line-source approximation, which assumes that membrane current is uniformly distributed within individual computational compartments and the medium is homogenous and isotropic^82^. Each model was simulated at a sampling rate at 30 kHz, identical to the acquisition rate of *in vivo* recordings. Each cell model received Poisson-like synaptic input (simulation time: 3s). We recorded the extracellular potential in a Neuropixels-like electrode array, which is a dense grid (5 µm spacing) consisting of 16 columns and 240 rows (total 3840 recording channels). To mimic Neuropixels recordings, we averaged extracellular potential within a 10 µm-by-10 µm area for each recording site. The extracellular action potential (EAP) was calculated based on the spike-triggered average of extracellular potentials.

### Data analysis

#### Feature extraction

Postprocessing included passing data through a 300 Hz high pass filter before extracting EAP waveforms. To classify cell types, we first extracted features from the extracellular waveform. With high density electrodes, we can record extracellular waveforms of a single unit from multiple sites. The recording site with largest amplitude (amplitude is the magnitude of the extremum of the waveform; **Fig. 2b**, *left*) is defined as the site closest to neuron soma, and the extracellular waveform recorded at this site we define as the one-channel waveform. Since the Neuropixels probe has four staggered columns of sites, we selected the two columns on the side of the probe with the largest one-channel amplitude for the one-as well as the multi-channel waveforms. The distance between sites is approximated by their vertical spacing (20 µm). The multi-channel waveform of a single unit includes EAPs from the channel with the largest EAP-amplitude and 10 additional channels above and below that location, spanning ±200 μm. Similarly, in the models, we selected the column of electrodes with the largest amplitude one-channel waveform. As expected, the channel with the largest EAP-amplitude in the models was located close to the soma and AIS location.

For the one-channel waveform (**Fig. 2b**, *left*), we calculated two features: TPW (trough-to-peak width) and REP (repolarization time). TPW measures the time that elapses from EAP trough (the global minimum of the curve) to EAP peak (the following local maximum). REP measures the time elapsed from EAP peak to the half of the peak value. These two EAP features capture different aspects of the intracellular potential, the speed of depolarization and of the subsequent after-hyperpolarization^17, 31^ and are commonly used to separate between fast-spiking (FS) units and regular-spiking (RS) units.

For the multi-channel waveform (**Fig. 2b**, *middle*), we extracted two additional features in the space domain: the inverse of the EAP propagation velocity below (1/*V_below_*) and above (1/*V_above_*) soma along the Neuropixels probe. Velocity measures how fast the EAP propagates along the probe with the point of reference being the EAP trough. When the EAP propagates fast, the time difference between two adjacent sites can sometimes be estimated as zero – to avoid infinite numbers, we calculated the inverse of velocity instead of velocity. A low value of inverse of velocity, indicates fast propagation. The inverse of propagation velocity below (1/*V_below_*) and above (1/*V_above_*) soma was then estimated by linear regression of the EAP trough at different sites against the distance of the sites relative to soma. We also define the spread of a unit by the range with amplitude above 12% of the maximum amplitude along the probe. Spread measures how far the waveform can propagate along a probe.

#### Symmetry index of EAP propagation

From the multi-channel EAP recordings, we defined a measure looking at the symmetry of spike propagation in the vertical direction above and below the spike initiation zone. Specifically, we defined the symmetry index (SI) as the distance between each point (1/*V_below_*, 1/*V_above_*) and the diagonal line (*y* = -*x*). The distance from point (*x0*, *y0*) to the line *ax + by + c =* 0 can be calculated by the following equations:

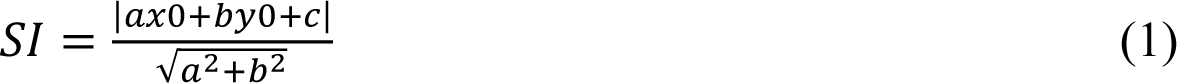

where (*x0*, *y0*) = (1/*V_below_*, 1/*V_above_*), and *a* = 1, *b* = 1, *c* = 0 for *y* = -*x*. A smaller value in the symmetry index indicates symmetric EAP propagation, while a larger value in the symmetry index indicates more asymmetric propagation.

#### Morphology bifurcation distance

The bifurcation distance (**w**) for one bifurcation node is defined as the projection of the line (**v**) from soma (S) to the position of the bifurcation node (N) projected to a line (**u**) connecting the soma (S) to a node (L) in y axis (**Fig. 5f**):

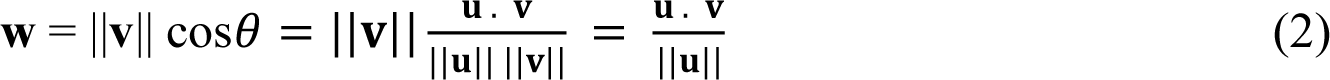

where 𝜃 is the angle between **u** and 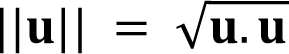 represents the length of the line **u**. The bifurcation distance is then normalized by the maximal absolute bifurcation distance for each neuron. We excluded the absolute bifurcation distance larger than 200 µm in the analysis because the node is too far away from the soma. The bifurcation distances above and below soma were calculated by the summation of the bifurcation distance for all the bifurcation nodes above and below soma, respectively. The sign of the bifurcation distances indicates the location of bifurcation nodes, where the positive sign indicates above soma, and negative sign indicates below soma. While comparing the bifurcation distances below vs. above the soma, we used the absolute value of the bifurcation distances.

#### Identification of EAP waveform clusters using K-means clustering

To identify cell clusters, we applied *K*-means clustering on the EAP features. *K*-means clustering is an unsupervised technique that seeks to find centroids that minimize the average Euclidian distance between points in the same cluster to the centroid. The optimal number of clusters was identified by two methods as in^22^.

One method is the standard elbow method that consists of plotting the within cluster sum of squares (WCSS) as a function of the number of clusters and picking the elbow of the curve as the number of optimal clusters. The global impact of all clusters’ distortions is given by the quantity:

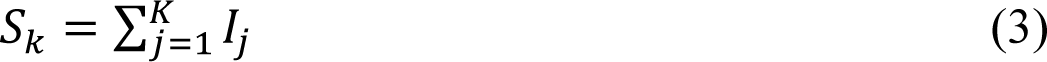

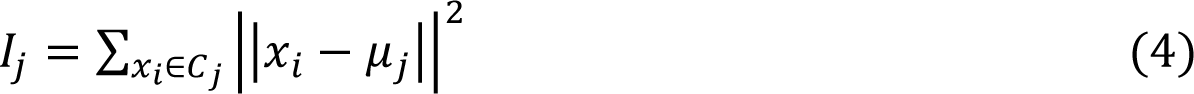

where 𝐼_𝑗_ is the distortion of cluster j that is a measure of the distance between points 𝑥_𝑖_ in cluster 𝐶_𝑗_ and its centroid 𝜇_𝑗_. In this paper, we have plotted WCSS curve as *S_k_* normalized by *S_1_*.

We also used a second method, the density function *f(K)*, that consists of plotting *f(K)* as a function of number of clusters and picking the minimal of the curve as the number of optimal clusters. The *f(K)* is from^83^:

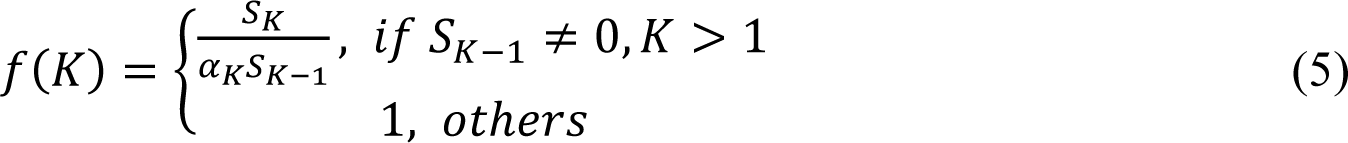

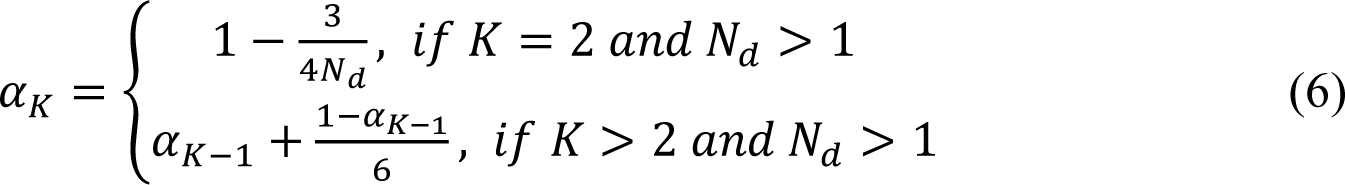

The value of *f(K)* is the ratio of the real distortion to the estimated distortion and is close to 1 when the data distribution is uniform. The smaller *f(K)*, the more concentrated the distribution.

We selected *K* based on these two methods and applied *K*-means to data with appropriate number of *K* for 1000 times with random initial values.

For the one-channel clustering, we used the standard one-channel waveform features (TPW and REP). To implement multi-channel clustering, we adopt the two one-channel clusters (RS and FS) and cluster each of them individually using the multi-channel features (1/*V_below_* and 1/*V_above_*).

#### Supervised machine learning for classification

The primary motivation for constructing the two-way classifiers was to bi-directional mapping between the experiment-based and model-based results. We built the experiment-based classifiers using on experimental EAP features and labels, then applied it to the model data to identify model neurons in the experimental space. Similarly, we built the model-based classifiers using model EAP features and labels, then applied it to the experimental data to identify experimental units in the model space. To train the classifier for the unbalanced FS and RS, before training, we have upsampled the ratio of FS and RS be 1:1. All classifications were performed with Monte-Carlo cross-validation consisting of a 100 “bootstrap composites” of individual classifiers (the partitions are done independently for each run) where the classifier was trained on a subset of the data (75%) and then the confusion matrix and accuracy were calculated on the left-out data (25%). We assigned the label based on the most frequently predicted label of the composite classifiers. For the classifier, we used a support vector machine (SVM) with a linear kernel (regularization parameter *C*=1) for two classes, or a random forest (gini criterion for splitting the nodes of a decision tree) for more than two classes.

#### Single unit firing properties

For this analysis, we only accounted for units with an EAP amplitude larger than 50 µV and a minimum of 100 spikes. Firing rate was calculated by *N/T* during the recording session, where N is the number of spikes and *T* is the total time in seconds. Coefficient of variation (*CV*) was calculated as the standard deviation of the interspike interval (ISI) divided by mean of ISI. The local variation (*LV*) is similar to *CV* but measures variation in adjacent ISIs and was calculated by^84^:

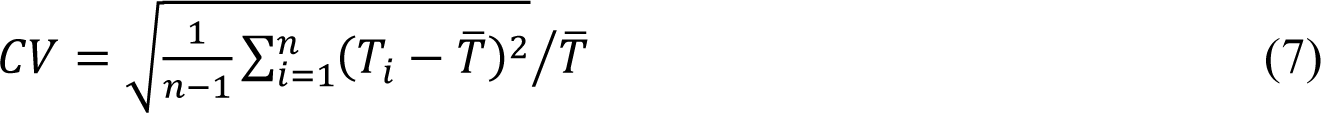

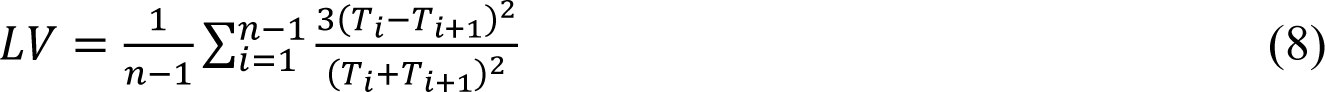

where 𝑇*_i_* is the duration of the *i*th ISI, n is the number of ISIs, and 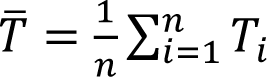 is the mean ISI.

#### Visual stimulus metrics

The three relevant visual stimulus metrics for drifting gratings used in the paper are f1/f0, the modulation index, and lifetime sparseness (**Table S2**).

##### f1/f0

the ratio of the 1st harmonic (response at the drifting frequency) to the 0^th^ harmonic (mean response). A high f1/f0 ratio indicates that the firing of the unit is modulated at the temporal frequency of the grating, while a low f1/f0 indicates that the unit fires relatively constantly during the presentation of the grating.

##### Modulation index (MI)

quantifies the phase-dependent responses to drifting gratings. MI measures the difference in power of the visually evoked response at a unit’s preferred stimulus frequency versus the average power spectrum^85^. MI > 3 corresponds to strong modulation of spiking at the stimulus frequency (indicative of simple-cell-like responses).

##### Lifetime sparseness

the selectivity of individual neurons to drifting gratings at different orientations and temporary frequencies was measured using lifetime sparseness, which captures the sharpness of a neuron’s mean response across different stimulus conditions^86^. A neuron that responds strongly to only a few conditions will have a lifetime sparseness close to 1 whereas a neuron that responds broadly to many conditions will have a lower lifetime sparseness. Detailed information about each metric is available at: https://allensdk.readthedocs.io/en/latest/visual_coding_neuropixels.html

##### Phase-locking Analysis

For the phase-locking analysis, we only include units with an EAP amplitude larger than 50 µV and a minimum of 100 spikes. For each unit, the maximal number of spikes considered in the analysis is limited to 10000. The percentage of phase-locked units was calculated by the number of units that fires at a preferred direction (assessed by the Rayleigh test) divided by the total number of units. To test whether spikes preferred certain phases of the LFP, the instantaneous phase of the LFP at several frequency bands (theta = 3-8 Hz, alpha = 8-12 Hz, beta = 12-30 Hz, low gamma = 30-50 Hz, high gamma = 50-90 Hz) was first calculated, using the Hilbert transform on each filtered LFP. 180^0^ is marked as the trough of the cycle. We chose pairs of units and LFPs recorded on different neighboring electrodes. Each spike was assigned with an instantaneous phase for each frequency band. A strongly phase-locked unit has a preferred direction in the phase histogram, while a weak phase-locked unit has no preferred direction in the phase histogram (**Fig. S6b**). To determine if a neuron exhibited a significant phase preference, we applied the Rayleigh test for non-uniformity. With the Rayleigh test, the null hypothesis is uniformity (e.g., no preferred direction), whereas the alternative is unimodality (e.g., a single preferred direction). A cell was considered phase-locked at a specific frequency range if the null hypothesis of uniformity of the phase distribution could be rejected at a p-value < 0.001 using a Rayleigh test^87, 88^. When the test indicated non-uniformity, the phase distribution was fitted to a circular normal distribution (von Mises distribution), with the concentration parameter (kappa) indicating the depth of the phase-locking in the direction of the mean phase. The inverse of kappa is analogous to variance of the normal distribution. For large kappa, the distribution becomes very concentrated around the mean phase, indicating a high phase-locking. Kappa values range from 0 to 1. Kappa, and preferred phase were calculated by a circular statistics toolbox pycircstat (https://github.com/circstat/pycircstat).

##### Detection of opto-tagged neurons

The peri-stimulus time histogram (PSTH) of spikes was used to present the light evoked neuronal responses. Time bins of 1 ms of PSTHs were used to measure the response to the light stimulation (square-wave pluses lasting 10ms). To prevent contamination by light artifacts, we only counted spikes in the window from 2 to 8 ms of the 10 ms light stimulation. The opto-tagged neuron was detected when the average firing rate across trials in the response window was higher than 25 Hz, and 2.5 times greater than its firing rate in a corresponding time window immediately preceding stimulus onset.

##### Statistical analysis

The Shapiro-Wilk test was used to determine whether the sample data have come from a normal distribution. The two-sample *t*-test (for normal distribution) or the nonparametric Mann–Whitney U test (for non-normal distribution) was used for statistical analysis of differences between means from two samples when appropriate. One-way ANOVA (for normal distribution) or the nonparametric Kruskal-Wallis H-test (for non-normal distribution) was used for comparisons across the multiple groups, with p-values corrected using the Holm-Bonferroni method (a step-down method using Bonferroni adjustments) for multiple tests. We used two sample z test for proportions to compare the percentages of phase locked cells between FS and RS and corrected the p-values via the Holm-Bonferroni method for multiple tests.

## Data Availability

The *in vivo* Neuropixels dataset is available for download in Neurodata Without Borders (NWB) format via the AllenSDK^23^: https://allensdk.readthedocs.io/en/latest/visual_coding_neuropixels.html

The Neurodata Without Borders files are also available on the DANDI Archive^23^: https://gui.dandiarchive.org/#/dandiset/000021

The *in vitro* electrophysiology data and the reconstructed morphology used to generate single-cell models are available in: https://celltypes.brain-map.org

The cell ID used in the paper was listed in the **Table S1**.

The optotagging experimental data set with Pvalb and Sst neurons is available through: https://allensdk.readthedocs.io/en/latest/_static/examples/nb/ecephys_optotagging.html

Source data are provided with this paper.

## Code Availability

The codes for calculating EAP features and clustering cell classes were custom written in Python and are made available on GitHub (https://github.com/yinawei/Mouse_V1_EAP_Analysis) with DOI (10.5281/zenodo.7679748).

The all-active mouse single-neuron models were generated using a Python pipeline and are also available on GitHub (https://github.com/yinawei/Mouse-all-active-models-EAP) with DOI (10.5281/zenodo.7679762).

## Acknowledgements

CAA thanks the Board of Governors of Cedars-Sinai Medical Center. CAA and SYL acknowledge the NIH grant RO1 NS120300-01. YW acknowledges the grants from National Natural Science Foundation of China (12101570) and the Scientific Project of Zhejiang Lab (2021KE0PI03, 2022KI0AC01, 2022KI0AC02). We thank the Allen Institute founder, Paul G. Allen.

## Author Contributions Statement

C.A.A. conceptualized this work. Y.W. and C.A.A. performed the research. A.N., W.V.G., and C.A.A. wrote the software to develop the all-active models. Y.W. wrote the analysis codes. X.J., J.H.S, D.D. and S.O. contributed the Neuropixels data. Y.W., A.N., and C.A.A. wrote the manuscript with input from all of the other authors (A.B., C.P.M., S.Y.L., X.J., J.H.S.).

## References

1. Gouwens, N. W. et al. Classification of electrophysiological and morphological neuron types in the mouse visual cortex. Nat. Neurosci. 22, 1182–1195 (2019).

2. Gouwens, N. W. et al. Integrated Morphoelectric and Transcriptomic Classification of Cortical GABAergic Cells. Cell 183, 935–953.e19 (2020).

3. Tasic, B. et al. Adult mouse cortical cell taxonomy revealed by single cell transcriptomics. Nat Neurosci vol. 19 335–46 (2016).

4. Tasic, B. et al. Shared and distinct transcriptomic cell types across neocortical areas. Nature 563, 72–78 (2018).

5. Harris, K. D. et al. Classes and continua of hippocampal CA1 inhibitory neurons revealed by single-cell transcriptomics. PLoS Biol 16, e2006387 (2018).

6. Tang, F. et al. mRNA-Seq whole-transcriptome analysis of a single cell. Nat Methods vol. 6 377–82 (2009).

7. Gupta, A., Wang, Y. & Markram, H. Organizing principles for a diversity of GABAergic interneurons and synapses in the neocortex. Science 287, 273–8 (2000).

8. Markram, H. et al. Interneurons of the neocortical inhibitory system. Nat Rev Neurosci 5, 793–807 (2004).

9. Cadwell, C. R. et al. Multimodal profiling of single-cell morphology, electrophysiology, and gene expression using Patch-seq. Nat Protoc 12, 2531–2553 (2017).

10. Jiang, X. et al. Principles of connectivity among morphologically defined cell types in adult neocortex. Science 350, aac9462 (2015).

11. Berens, P. et al. Community-based benchmarking improves spike rate inference from two-photon calcium imaging data. PLoS Comput Biol 14, e1006157 (2018).

12. Huang, L. et al. Relationship between simultaneously recorded spiking activity and fluorescence signal in GCaMP6 transgenic mice. Elife vol. 10 (2021).

13. de Vries, S. E. J. et al. A large-scale standardized physiological survey reveals functional organization of the mouse visual cortex. Nat Neurosci 23, 138–151 (2020).

14. Bartho, P. et al. Characterization of neocortical principal cells and interneurons by network interactions and extracellular features. J Neurophysiol 92, 600–8 (2004).

15. Peyrache, A. & Destexhe, A. Electrophysiological monitoring of inhibition in mammalian species, from rodents to humans. Neurobiol Dis vol. 130 104500 (2019).

16. Peyrache, A. et al. Spatiotemporal dynamics of neocortical excitation and inhibition during human sleep. Proc Natl Acad Sci U S A vol. 109 1731–6 (2012).

17. Trainito, C., von Nicolai, C., Miller, E. K. & Siegel, M. Extracellular Spike Waveform Dissociates Four Functionally Distinct Cell Classes in Primate Cortex. Curr Biol vol. 29 2973–2982.e5 (2019).

18. McCormick, D. A., Connors, B. W., Lighthall, J. W. & Prince, D. A. Comparative electrophysiology of pyramidal and sparsely spiny stellate neurons of the neocortex. J Neurophysiol 54, 782–806 (1985).

19. Ardid, S. et al. Mapping of functionally characterized cell classes onto canonical circuit operations in primate prefrontal cortex. J Neurosci vol. 35 2975–91 (2015).

20. Katai, S. et al. Classification of extracellularly recorded neurons by their discharge patterns and their correlates with intracellularly identified neuronal types in the frontal cortex of behaving monkeys. Eur J Neurosci vol. 31 1322–38 (2010).

21. Jun, J. J. et al. Fully integrated silicon probes for high-density recording of neural activity. Nature vol. 551 232–236 (2017).

22. Jia, X. et al. High-density extracellular probes reveal dendritic backpropagation and facilitate neuron classification. J Neurophysiol vol. 121 1831–1847 (2019).

23. Siegle, J. H. et al. Survey of spiking in the mouse visual system reveals functional hierarchy. Nature (2021).

24. Nandi, A., et al. Single-neuron models linking electrophysiology, morphology and transcriptomics across cortical cell types. bioRxiv 2020.04.09.030239 (2020).

25. Anastassiou, C. A., Perin, R., Buzsáki, G., Markram, H. & Koch, C. Cell type- and activity-dependent extracellular correlates of intracellular spiking. J Neurophysiol vol. 114 608–23 (2015).

26. Gold, C., Henze, D. A., Koch, C. & Buzsáki, G. On the origin of the extracellular action potential waveform: A modeling study. J Neurophysiol 95, 3113–28 (2006).

27. Mosher, C. P. et al. Cellular Classes in the Human Brain Revealed In Vivo by Heartbeat-Related Modulation of the Extracellular Action Potential Waveform. Cell Rep. 30, 3536–3551.e6 (2020).

28. Buzsáki, G., Anastassiou, C. A. & Koch, C. The origin of extracellular fields and currents--EEG, ECoG, LFP and spikes. Nat. Rev. Neurosci. 13, 407–420 (2012).

29. Nicholson, C. & Freeman, J. A. Theory of current source-density analysis and determination of conductivity tensor for anuran cerebellum. J. Neurophysiol. 38, 356–368 (1975).

30. Buchin, A., et al. Multi-modal characterization and simulation of human epileptic circuitry. bioRxiv 2020.04.24.060178 (2020).

31. Henze, D. A. et al. Intracellular features predicted by extracellular recordings in the hippocampus in vivo. J Neurophysiol 84, 390–400 (2000).

32. Buzsáki, G. & Kandel, A. Somadendritic backpropagation of action potentials in cortical pyramidal cells of the awake rat. J. Neurophysiol. 79, 1587–1591 (1998).

33. Chen, G. et al. Distinct Inhibitory Circuits Orchestrate Cortical beta and gamma Band Oscillations. Neuron 96, 1403–1418.e6 (2017).

34. Senzai, Y., Fernandez-Ruiz, A. & Buzsáki, G. Layer-Specific Physiological Features and Interlaminar Interactions in the Primary Visual Cortex of the Mouse. Neuron 101, 500–513.e5 (2019).

35. Quyen, M. L. V. et al. High-frequency oscillations in human and monkey neocortex during the wake–sleep cycle. Proc. Natl. Acad. Sci. 113, 9363–9368 (2016).

36. Csicsvari, J., Hirase, H., Czurkó, A., Mamiya, A. & Buzsáki, G. Oscillatory coupling of hippocampal pyramidal cells and interneurons in the behaving rat. J. Neurosci. 19, 274–287 (1999).

37. Perez-Reyes, E. Molecular Physiology of Low-Voltage-Activated T-type Calcium Channels. Physiol. Rev. 83, 117–161 (2003).

38. Tremblay, R., Lee, S. & Rudy, B. GABAergic Interneurons in the Neocortex: From Cellular Properties to Circuits. Neuron 91, 260–292 (2016).

39. Erisir, A., Lau, D., Rudy, B. & Leonard, C. S. Function of Specific K ^+^ Channels in Sustained High-Frequency Firing of Fast-Spiking Neocortical Interneurons. J. Neurophysiol. 82, 2476– 2489 (1999).

40. Lien, C.-C. & Jonas, P. Kv3 Potassium Conductance is Necessary and Kinetically Optimized for High-Frequency Action Potential Generation in Hippocampal Interneurons. J. Neurosci. 23, 2058–2068 (2003).

41. McMahon, A. et al. Allele-dependent changes of olivocerebellar circuit properties in the absence of the voltage-gated potassium channels Kv3.1 and Kv3.3: Olivocerebellar system properties in the absence of Kv3 channels. Eur. J. Neurosci. 19, 3317–3327 (2004).

42. Pettersen, K. H. & Einevoll, G. T. Amplitude variability and extracellular low-pass filtering of neuronal spikes. Biophys. J. 94, 784–802 (2008).

43. Kvitsiani, D. et al. Distinct behavioural and network correlates of two interneuron types in prefrontal cortex. Nature vol. 498 363–6 (2013).

44. Lima, S. Q., Hromadka, T., Znamenskiy, P. & Zador, A. M. PINP: a new method of tagging neuronal populations for identification during in vivo electrophysiological recording. PLoS One vol. 4 e6099 (2009).

45. Jia, X., et al. Multi-area functional modules mediate feedforward and recurrent processing in visual cortical hierarchy. bioRxiv 2020.08.30.272948 (2020).

46. Steinmetz, N. A., Zatka-Haas, P., Carandini, M. & Harris, K. D. Distributed coding of choice, action and engagement across the mouse brain. Nature 576, 266–273 (2019).

47. Gouwens, N. W. et al. Classification of electrophysiological and morphological neuron types in the mouse visual cortex. Nat. Neurosci. 22, 1182–1195 (2019).

48. Sakata, S. & Harris, K. Laminar-dependent effects of cortical state on auditory cortical spontaneous activity. Front. Neural Circuits 6, (2012).

49. Petersen, C. C. H. & Crochet, S. Synaptic Computation and Sensory Processing in Neocortical Layer 2/3. Neuron 78, 28–48 (2013).

50. Senzai, Y., Fernandez-Ruiz, A. & Buzsáki, G. Layer-Specific Physiological Features and Interlaminar Interactions in the Primary Visual Cortex of the Mouse. Neuron 101, 500–513.e5 (2019).

51. Jia, X. et al. Multi-regional module-based signal transmission in mouse visual cortex. Neuron 110, 1585–1598.e9 (2022).

52. Siegle, J. H. et al. Survey of spiking in the mouse visual system reveals functional hierarchy. Nature 1–7 (2021) doi:10.1038/s41586-020-03171-x.

53. de Vries, S. E. J. et al. A large-scale standardized physiological survey reveals functional organization of the mouse visual cortex. Nat. Neurosci. 23, 138–151 (2020).

54. Freund, T. f. & Buzsáki, G. Interneurons of the hippocampus. Hippocampus 6, 347–470 (1996).

55. Klausberger, T. et al. Brain-state-and cell-type-specific firing of hippocampal interneurons in vivo. Nature 421, 844–848 (2003).

56. Lapray, D. et al. Behavior-dependent specialization of identified hippocampal interneurons. Nat. Neurosci. 15, 1265–1271 (2012).

57. Einstein, M. C., Polack, P.-O., Tran, D. T. & Golshani, P. Visually Evoked 3–5 Hz Membrane Potential Oscillations Reduce the Responsiveness of Visual Cortex Neurons in Awake Behaving Mice. J. Neurosci. 37, 5084–5098 (2017).

58. Lasztóczi, B. & Klausberger, T. Layer-Specific GABAergic Control of Distinct Gamma Oscillations in the CA1 Hippocampus. Neuron 81, 1126–1139 (2014).

59. Schomburg, E. W. Biophysical and network mechanisms of high frequency extracellular potentials in the rat hippocampus. (California Institute of Technology, 2014).

60. Tukker, J. J., Fuentealba, P., Hartwich, K., Somogyi, P. & Klausberger, T. Cell Type-Specific Tuning of Hippocampal Interneuron Firing during Gamma Oscillations In Vivo. J. Neurosci. 27, 8184–8189 (2007).

61. Bannister, N. J. & Larkman, A. U. Dendritic morphology of CA1 pyramidal neurones from the rat hippocampus: I. Branching patterns. J. Comp. Neurol. 360, 150–160 (1995).

62. Bannister, N. J. & Larkman, A. U. Dendritic morphology of CA1 pyramidal neurones from the rat hippocampus: II. Spine distributions. J. Comp. Neurol. 360, 161–171 (1995).

63. Cembrowski, M. S. et al. Spatial Gene-Expression Gradients Underlie Prominent Heterogeneity of CA1 Pyramidal Neurons. Neuron 89, 351–368 (2016).

64. Deguchi, Y., Donato, F., Galimberti, I., Cabuy, E. & Caroni, P. Temporally matched subpopulations of selectively interconnected principal neurons in the hippocampus. Nat. Neurosci. 14, 495–504 (2011).

65. Hodge, R. D. et al. Conserved cell types with divergent features in human versus mouse cortex. Nature 1–8 (2019) doi:10.1038/s41586-019-1506-7.

66. Lee, S.-H. et al. Parvalbumin-Positive Basket Cells Differentiate among Hippocampal Pyramidal Cells. Neuron 82, 1129–1144 (2014).

67. Soltesz, I. & Losonczy, A. CA1 pyramidal cell diversity enabling parallel information processing in the hippocampus. Nat. Neurosci. 21, 484–493 (2018).

68. Yao, Z. et al. A taxonomy of transcriptomic cell types across the isocortex and hippocampal formation. Cell 184, 3222–3241.e26 (2021).

69. Berg, J. et al. Human neocortical expansion involves glutamatergic neuron diversification. Nature 598, 151–158 (2021).

70. Graves, A. R. et al. Hippocampal Pyramidal Neurons Comprise Two Distinct Cell Types that Are Countermodulated by Metabotropic Receptors. Neuron 76, 776–789 (2012).

71. Mizuseki, K., Diba, K., Pastalkova, E. & Buzsáki, G. Hippocampal CA1 pyramidal cells form functionally distinct sublayers. Nat. Neurosci. 14, 1174–1181 (2011).

72. Destexhe, A., Rudolph, M. & Paré, D. The high-conductance state of neocortical neurons in vivo. Nat. Rev. Neurosci. 4, 739–751 (2003).

73. Steriade, M., Timofeev, I. & Grenier, F. Natural Waking and Sleep States: A View From Inside Neocortical Neurons. J. Neurophysiol. 85, 1969–1985 (2001).

74. Timofeev, I., Grenier, F., Bazhenov, M., Sejnowski, T. J. & Steriade, M. Origin of Slow Cortical Oscillations in Deafferented Cortical Slabs. Cereb. Cortex 10, 1185–1199 (2000).

75. Timofeev, I., Grenier, F. & Steriade, M. Disfacilitation and active inhibition in the neocortex during the natural sleep-wake cycle: An intracellular study. Proc. Natl. Acad. Sci. 98, 1924– 1929 (2001).

76. Kalmbach, B. E. et al. h-Channels Contribute to Divergent Intrinsic Membrane Properties of Supragranular Pyramidal Neurons in Human versus Mouse Cerebral Cortex. Neuron 100, 1194–1208.e5 (2018).

77. Hu, H., Martina, M. & Jonas, P. Dendritic Mechanisms Underlying Rapid Synaptic Activation of Fast-Spiking Hippocampal Interneurons. Science 327, 52–58 (2010).

78. Stuart, G., Spruston, N., Sakmann, B. & Hausser, M. Action potential initiation and backpropagation in neurons of the mammalian CNS. Trends Neurosci 20, 125–31 (1997).

79. Madisen, L. et al. A toolbox of Cre-dependent optogenetic transgenic mice for light-induced activation and silencing. Nat. Neurosci. 15, 793–802 (2012).

80. Zhang, F., Wang, L. P., Boyden, E. S. & Deisseroth, K. Channelrhodopsin-2 and optical control of excitable cells. Nat Methods 3, 785–92 (2006).

81. Stringer, C. et al. Spontaneous behaviors drive multidimensional, brainwide activity. Science 364, 255 (2019).

82. Gratiy, S. L. et al. BioNet: A Python interface to NEURON for modeling large-scale networks. PLoS One vol. 13 e0201630 (2018).

83. Pham, D. T., Dimov, S. S. & Nguyen, C. D. Selection of K in K-means clustering. Proceedings of the Institution of Mechanical Engineers, Part C: Journal of Mechanical Engineering Science vol. 219 103–119 (2005).

84. Shinomoto, S., Shima, K. & Tanji, J. Differences in spiking patterns among cortical neurons. Neural Comput 15, 2823–42 (2003).

85. Matteucci, G., Bellacosa Marotti, R., Riggi, M., Rosselli, F. B. & Zoccolan, D. Nonlinear Processing of Shape Information in Rat Lateral Extrastriate Cortex. J Neurosci 39, 1649–1670 (2019).

86. Vinje, W. E. & Gallant, J. L. Sparse coding and decorrelation in primary visual cortex during natural vision. Science 287, 1273–6 (2000).

87. Fisher, N. I. Statistical analysis of circular data. (Cambridge University Press, 1993).

88. Jacobs, J., Kahana, M. J., Ekstrom, A. D. & Fried, I. Brain oscillations control timing of single-neuron activity in humans. J Neurosci 27, 3839–44 (2007).

