## Supplemental_Material for "Associations between *in vitro*, *in vivo* and *in silico* cell classes in mouse primary visual cortex"

### SUPPLEMENTARY INFORMATION

**Figure S1**

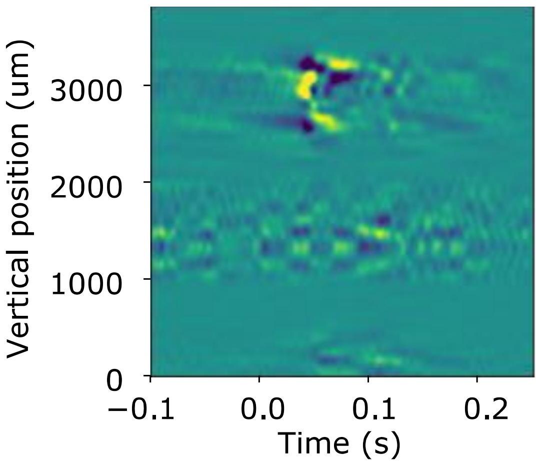


**Figure S1**. Visual stimulus (flashes) evoked a strong response evident in the current source density (CSD) analysis. X axis is the time relative to stimulus onset (0: time of stimulus onset). Y axis is the vertical position of the Neuropixels probe (associated with the cortical depth). The sink of CSD (yellow) first occurred at the vertical position of 2740 um. This position is ascribed as the middle of layer 4 and denoted as depth 0.

**Figure S2**

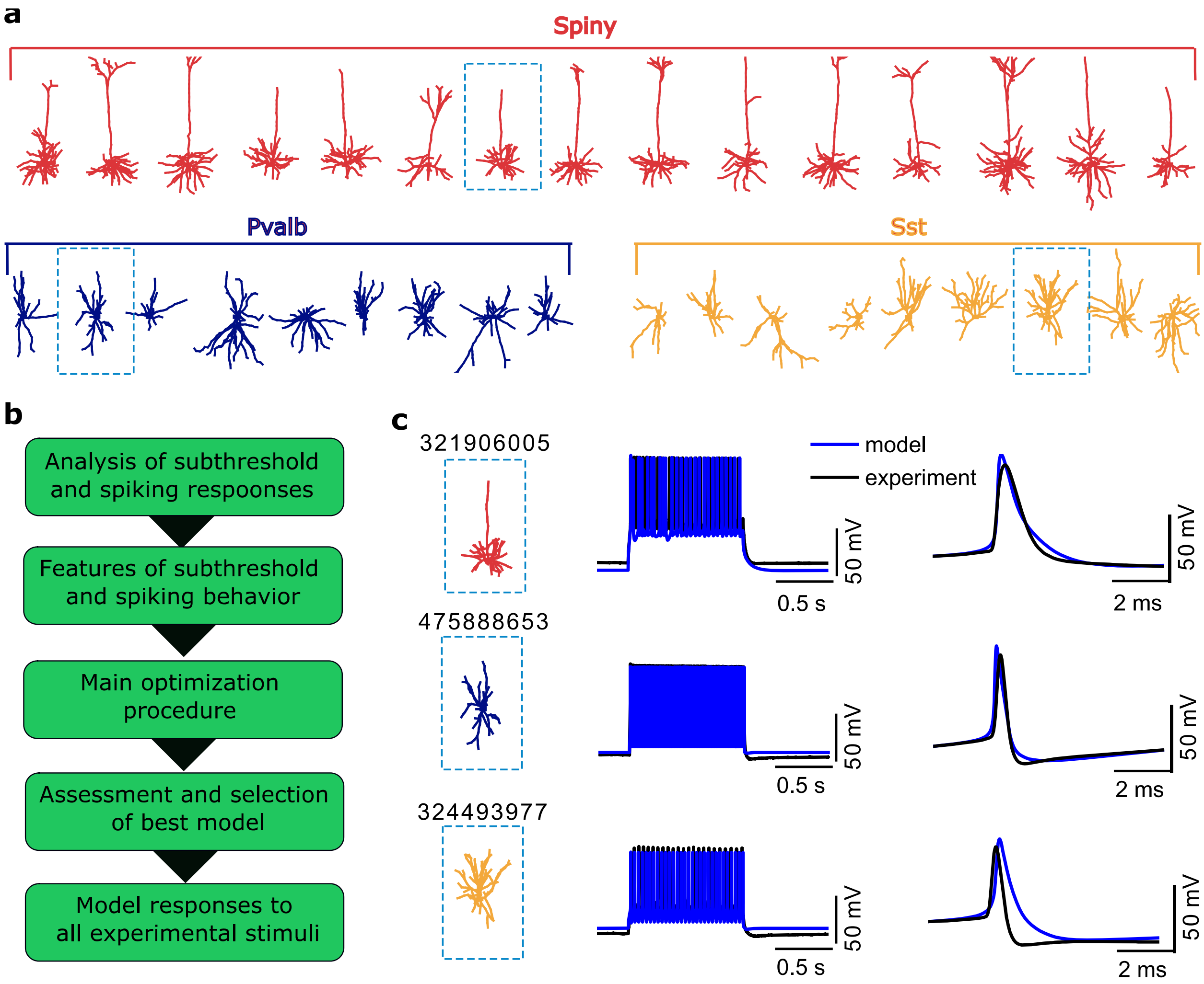


**Figure S2. Computational optimization workflow for the generation of single-cell ("all-active") models of mouse cortical neurons. a)** The morphology of spiny (red), and aspiny (blue: Pvalb; orange: Sst) neurons is reconstructed from imaged biocytin fills performed in whole-cell patch-clamp experiments. **b)** Sequence of steps comprising the model generation and optimization workflow. **c)** Three example cell models (top: a spiny cell; middle: a Pvalb cell; bottom: a Sst cell). Left column: reconstructed neuron morphology (box indicates the morphology in panel **a**); Middle column: simulated dynamics for the individual models resulting from the optimization workflow (intracellular stimulus: 1s somatic current injection; black: experimental trace; blue: model trace). Right column: the somatic action potential waveform. For more information on the computational optimization procedure see Methods.

**Figure S3**

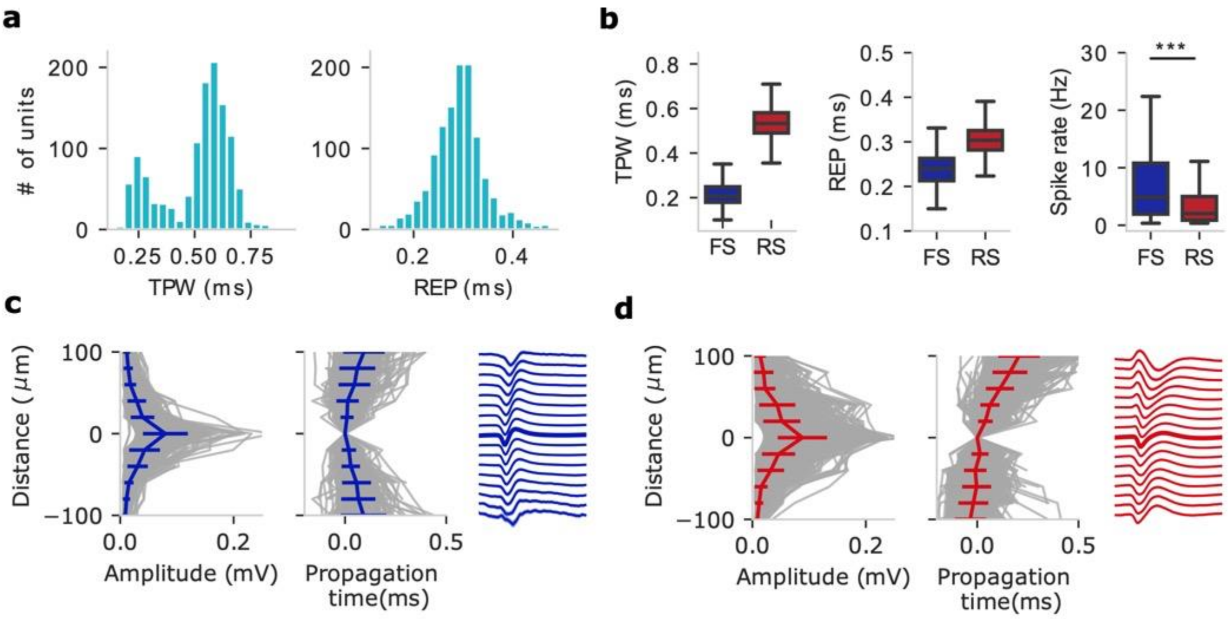


**Figure S3**. **One- and multi-channel extracellular action potential (EAP) feature analysis of one-channel clusters RS and FS. a**) The distribution of one-channel EAP features across all recorded units: trough-peak width (TPW) and repolarization time (REP). **b**) Comparison of the spike rate shows that the narrow TPW cluster constitutes a fast-spiking (FS) group while the wide TPW cluster is attributed to a regular-spiking (RS) group. Box plots show center line as median, box limits as upper (75%) and lower (25%) quartiles. The whiskers extend from the box limits by 1x the interquartile range. n=281 FS units, n=923 RS units. Mann-Whitney U test, two-sided, $p=3.55*10^{-18}$, ***$p<0.001$. **c-d**) EAP amplitude and propagation time (calculated as a multi-channel feature) of FS (**c,** n=281) and RS (**d**, n=923) as function of the cortical depth. Gray lines indicate individual units while colored lines indicate mean ± SD (standard deviation). Note the asymmetry in propagation time of EAPs between FS (**c**, blue) and RS (**d**, red).

**Figure S4**

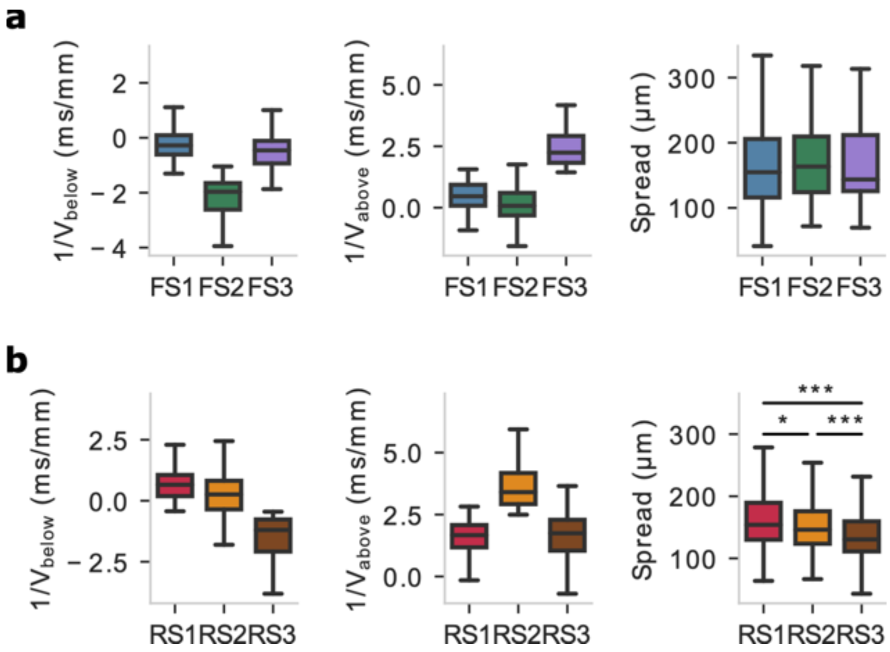


**Figure S4**. **Multi-channel extracellular action potential (EAP) feature analysis of multi-channel clusters RS1-3 and FS1-3.** Comparisons and boxplots of multi-channel extracellular action potential (EAP) features among (**a**) fasting-spiking (FS) clusters: FS1 (n=130), FS2 (n=82), FS3 (n=69), and (**b**) fasting-spiking (FS) clusters: RS1 (n=479), RS2 (n=235), RS3 (n=209) . Box plots show center line as median, box limits as upper and lower quartiles. The notches represent confidence intervals around the median. The whiskers extend from the box limits by 1x the interquartile range. Comparison of spread among FS1-3, Kruskal-Wallis H-test, F=0.457, p=0.795, p-values corrected using the Holm-Bonferroni method for multiple tests; Comparison of spread among RS1-3, Kruskal-Wallis H-test, F=48.2, p = $3.34*10^{-11}$, p-values corrected using the Holm-Bonferroni method for multiple tests, *p<0.05, ***p<0.001.

**Figure S5**

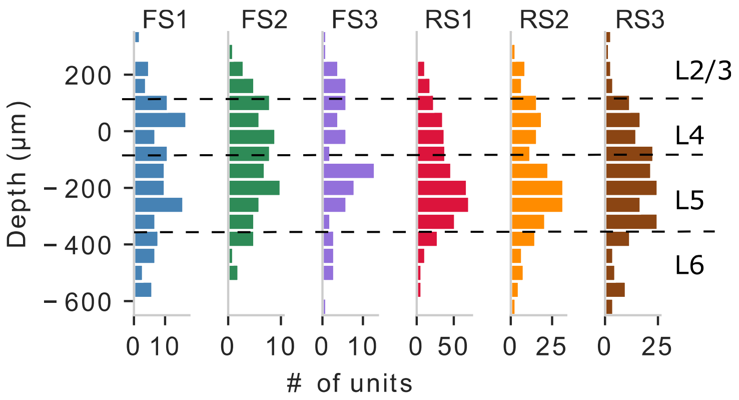


**Figure S5**. **Distribution of extracellular action potential (EAP) multi-channel waveform clusters along the cortical depth.** Distribution of EAP waveform clusters (FS1-3 and RS1-3) in mouse primary visual cortex (V1) as function of cortical depth relative to layer 4 (depth 0 indicates the center of layer 4). Localization of layer 4 center is shown in **Fig. S1**.

**Figure S6**

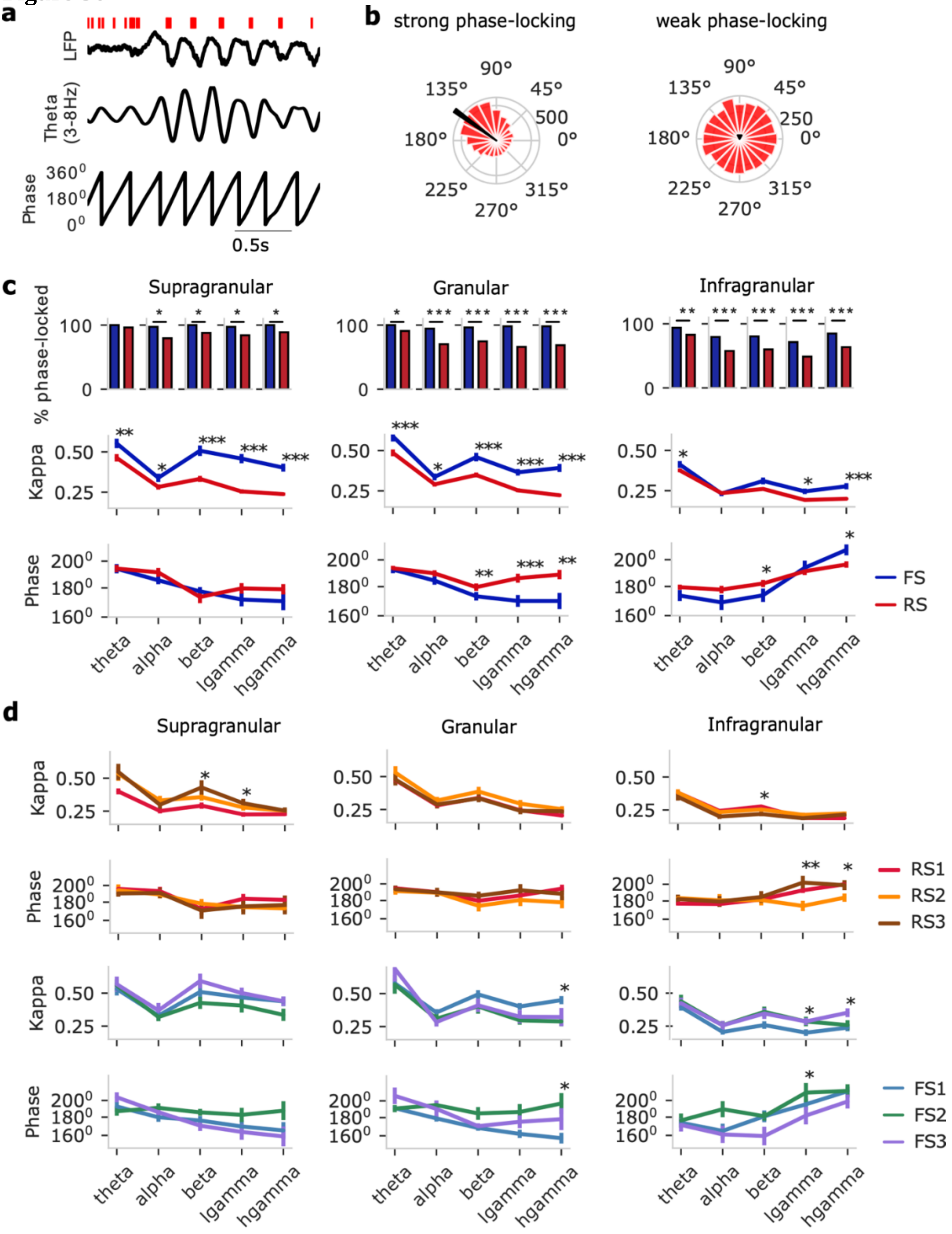


**Figure S6**. **Spike-phase relationships of one- and multi-channel clusters. a**) Example of raw local field potential (LFP, top, black trace) and the raster plot of a spiking unit recorded simultaneously (top, red), the filtered LFP at theta band (middle) and the instantaneous phase computed via the Hilbert transform (bottom). $180^0$ (or $\pi$) represent the trough of each cycle. **b**) Examples of the phase distribution of a strongly phase-locked unit (left), and a weakly entrained unit (right). The direction of the black arrow represents the mean preferred phase, while the length of the black arrow represents the kappa. **c**) Phase locking across layers. Top: The percentage of phase-locked units in fast-spiking (FS, blue) vs. regular-spiking (RS, red) at different LFP frequency bands: theta, alpha, beta, low gamma (lgamma), high gamma (hgamma). Supragranular layer: *n*=37 FS, *n*=107 RS; Granular layer: *n* = 54 FS, *n*=185 RS; Infragranular layer: *n*=112 FS, *n*=453 RS. Two sample z test for proportions with p-values corrected by the Holm-Bonferroni method for multiple tests. *p<0.05, **p<0.01, ***p<0.001. Bottom: kappa and preferred phase. Data are presented as mean ± SEM (standard error of mean) . Mann-Whitney U test, two-sided, *p<0.05, **p<0.01, ***p<0.001. **d**) Kappa and preferred phase of RS1-3 and FS1-3 across layers. Top: RS1 (red), RS2 (orange), and RS3 (brown); Bottom: FS1 (blue), FS2 (green), FS3 (purple). Data are presented as mean ± SEM. Mann-Whitney U test, two-sided, *p<0.05, **p<0.01.

**Figure S7**

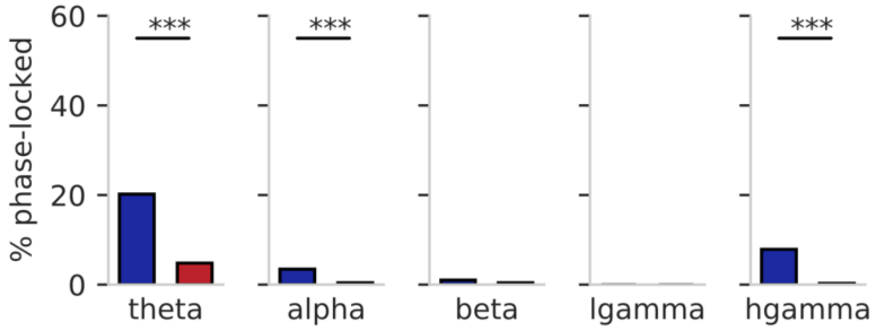


**Figure S7**. **Spike-phase relationship on one-channel clusters during spontaneous activity.** The percentage of phase-locked units during spontaneous activity for FS (fast-spiking, blue, n=203) and RS (regular-spiking, red, n=750) clusters at different LFP (local field potential) frequency bands: theta, alpha, beta, low gamma (lgamma), high gamma (hgamma). Two sample z test for proportions with p values corrected by the Holm-Bonferroni method for multiple tests. ***$p<0.001$. The total number of units activated during spontaneous activity are *n*=1705 but we only included the units activated during both spontaneous activity and drifting gratings (*n*=953) to show the input- and behavior-dependent properties.

Figure S8

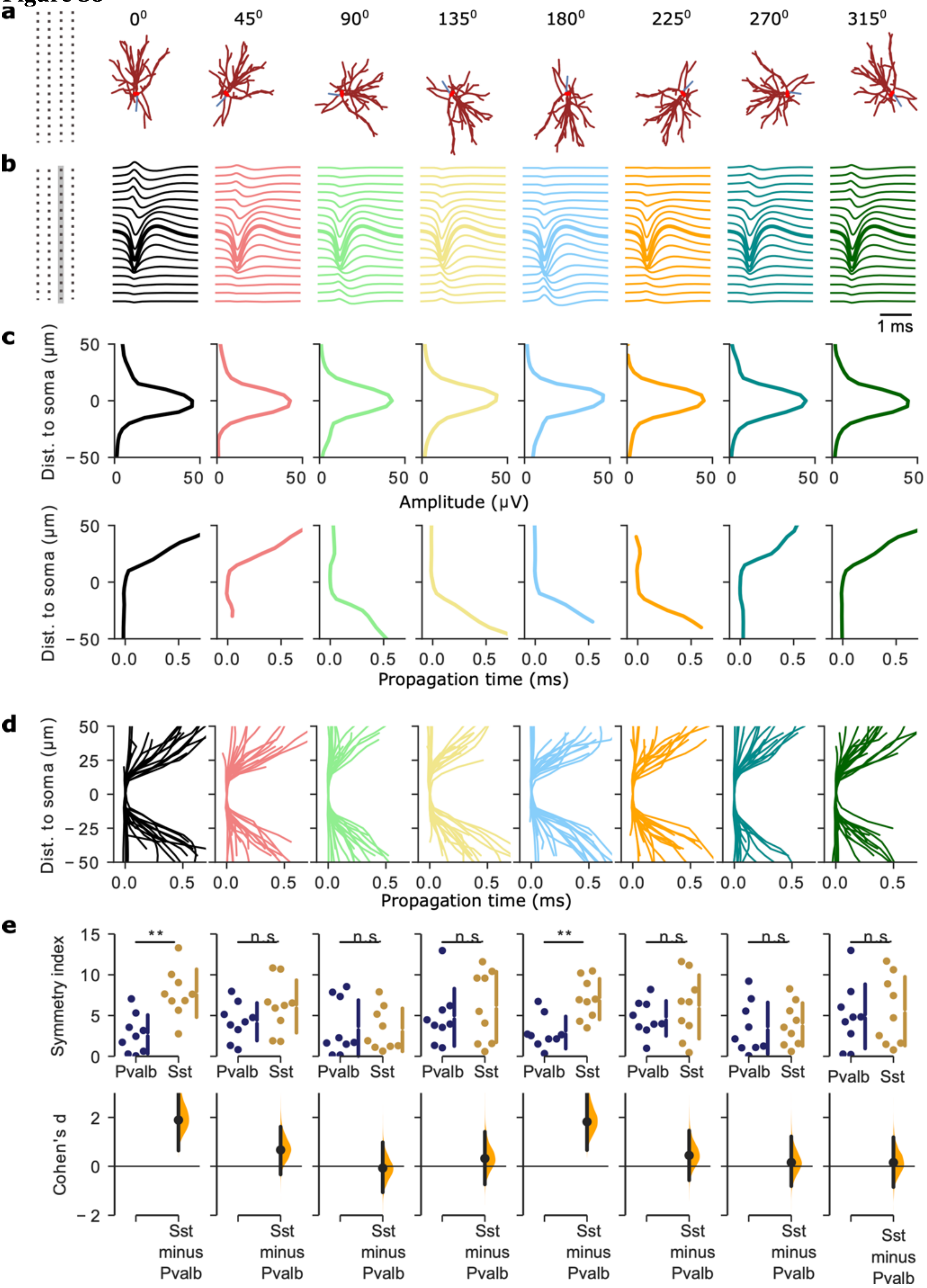


**Figure S8**. **The effect of the angle between cellular morphology and the simulated Neuropixels probe traversing along the cortical depth axis**. **a-c**) Example of the rotation effect of the cellular morphology on the extracellular action potential (EAP) features of one Sst cell (cell ID: 324493977). **b**) The multi-channel EAPs correspond to the rotation angles (shown in panel **a**). **c**) EAP features can change as a function of distance-to-soma at different rotation angles. The rotation effect is shown on two EAP features: amplitude (top) and propagation time (bottom). While the EAP amplitude remains broadly unchanged with rotation, the EAP propagation time changes considerably with rotation angle. **d-e**) The rotation effect on the EAP features of Pvalb and Sst is shown across cellular models (Pvalb: n = 9; Sst: n = 9; see also **Table S1**). **d**) The EAP propagation time is shown as a function of distance to soma for all aspiny models at different rotation angles (reported in panel **a**, top). Each line corresponds to one cell. **e)** Symmetry index of the EAP propagation changes with rotation angle with certain rotation angles exacerbating differences in symmetry index between Pvalb vs. Sst while others leave them statistically undetected (dark blue: Pvalb, n=9; orange: Sst, n=9). Top: symmetry index between Pvalb and Sst for various rotation angles (from left to right, as shown in panel **a**) and models. Each dot represents one model. Lines designate mean ± standard deviation (SD). Bottom: Cohen's d shows the effect size and direction. **p<0.01, n.s. represents no significance.

**Table S1. Metadata about origin of experimental *in vitro* data used to generate single-cell models**. *In vitro* electrophysiology data and the reconstructed morphology are generated for each whole-cell patched neuron from mouse primary visual cortex. Data for each experiment available online (https://celltypes.brain-map.org).

| | Cell ID | Cre lines | Layer | Dendrite type |
|---|---|---|---|---|
| 1 | 395830185 | Scnn1a | L4 | spiny |
| 2 | 476218657 | Scnn1a | L4 | spiny |
| 3 | 320207387 | Scnn1a | L4 | spiny |
| 4 | 422738880 | Nr5a1 | L4 | spiny |
| 5 | 476457450 | Nr5a1 | L4 | spiny |
| 6 | 386049446 | Nr5a1 | L4 | spiny |
| 7 | 321906005 | Nr5a1 | L4 | spiny |
| 8 | 485931158 | Rorb | L4 | spiny |
| 9 | 479179020 | Rorb | L4 | spiny |
| 10 | 478586295 | Rorb | L5 | spiny |
| 11 | 468120757 | Scnn1a | L5 | spiny |
| 12 | 484559000 | Rorb | L5 | spiny |
| 13 | 476048909 | Rorb | L5 | spiny |
| 14 | 314822529 | Rorb | L5 | spiny |
| 15 | 318808427 | Nr5a1 | L2/3 | spiny |
| 16 | 487667205 | Pvalb | L4 | aspiny |
| 17 | 475888653 | Pvalb | L4 | aspiny |
| 18 | 471085676 | Pvalb | L5 | aspiny |
| 19 | 469801569 | Pvalb | L5 | aspiny |
| 20 | 469610831 | Pvalb | L5 | aspiny |
| 21 | 396608557 | Pvalb | L5 | aspiny |
| 22 | 333604946 | Pvalb | L2/3 | aspiny |
| 23 | 330080937 | Pvalb | L5 | aspiny |
| 24 | 318331342 | Pvalb | L4 | aspiny |
| 25 | 485058595 | Sst | L5 | aspiny |
| 26 | 482690728 | Sst | L5 | aspiny |
| 27 | 478396248 | Sst | L5 | aspiny |
| 28 | 466664172 | Sst | L5 | aspiny |
| 29 | 485466109 | Sst | L2/3 | aspiny |
| 30 | 473540161 | Sst | L2/3 | aspiny |
| 31 | 324493977 | Sst | L2/3 | aspiny |
| 32 | 476104386 | Sst | L6 | aspiny |
| 33 | 313862306 | Sst | L6 | aspiny |
